# Fungal effector Jsi1 hijacks plant JA/ET signaling through Topless

**DOI:** 10.1101/844365

**Authors:** Martin Darino, Joana Marques, Khong-Sam Chia, David Aleksza, Luz Mayela Soto, Simon Uhse, Michael Borg, Ruben Betz, Janos Bindics, Krzysztof Zienkiewicz, Ivo Feussner, Yohann Petit-Houdenot, Armin Djamei

## Abstract

*Ustilago maydis (U. maydis)* is the causal agent of maize smut disease. During the colonization process, the fungus secretes effector proteins which suppress immune responses and redirect the host-metabolism in favor of the pathogen. Here we describe a novel strategy by which *U. maydis* induces plant jasmonate/ethylene (JA/ET) hormone signaling and thereby biotrophic susceptibility. The *U. maydis* effector *Jasmonate/Ethylene signaling inducer 1* (*Jsi1*) possesses an ethylene-responsive element binding factor-associated amphiphilic repression (EAR) motif, DLNxxP, which interacts with the second WD40 domain of the conserved plant co-repressor family Topless/Topless related (TPL/TPR). Jsi1 interaction with TPL/TPRs leads to derepression of the ethylene response factor (ERF) branch of the JA/ET signaling pathway, supporting biotrophic susceptibility. Jsi1 likely activates the ERF branch by interfering with the binding of endogenous DLNxxP motif-containing ERF transcription factors to TPL/TPR proteins. The identification of effector proteins possessing a DLNxxP motif in different fungal species with biotrophic and hemibiotrophic lifestyles together with the validation of the interaction between such effectors from *Magnaporthe oryzae* (*M. oryzae*), *Sporisorium scitamineum* (*S. scitamineum*), and *S. reilianum* with TPL proteins indicates the convergent evolution of effectors for modulating the TPL/TPR co-repressor hub.

## Introduction

The biotrophic fungus *U. maydis* causes smut disease, producing galls on all aerial parts of its host-plant maize. During colonization, the fungus secretes numerous manipulative molecules, termed effectors, which interfere with the host’s cellular machinery to suppress plant defense responses, redirect development, and enhance nutrient access (Win et al., 2012). As effectors play a critical role during plant colonization, their identification and functional characterization is essential to understand the processes involved in establishment of the plant-pathogen interaction. Effectors generally have very low sequence conservation, likely a result of the high co-evolutionary pressure to evade recognition by the host’s immune system while optimizing host manipulation. In *U. maydis*, 467 genes were classified as putative secreted proteins and 203 of these encode proteins of unknown function (Lanver et al., 2017). Only a few have been characterized as effector proteins, with diverse functions during the biotrophic phase. Three apoplastic effectors, Pep1, Pit2, and Rsp3, interfere with host peroxidases, cysteine proteases, and mannose binding proteins, respectively, thereby inhibiting early immune responses of the host plant (Ma et al., 2018, Mueller et al., 2013, Hemetsberger et al., 2012). Three translocated effectors, Tin2, Cmu1, and See1, have also been characterized. Tin2 induces anthocyanin formation to reduce cell wall lignification (Tanaka et al., 2014). Cmu1 interferes with the host chorismate metabolism, reducing the level of the plant defense hormone salicylic acid (SA) (Djamei et al., 2011). See1 alters DNA synthesis to promote gall formation (Redkar et al., 2015).

Plants coordinate pathogen-specific immune responses through an elaborate crosstalk between hormone signaling pathways. Activation of SA signaling usually leads to activation of immune responses against biotrophic and hemibiotrophic pathogens. In contrast, activation of jasmonate (JA) signaling leads to activation of immune responses to necrotrophic pathogens and insect herbivory. While ethylene (ET) signaling can be synergistic with JA signaling, SA and JA signaling are generally antagonistic to one another (Pieterse et al., 2012).

The JA signaling pathway is activated upon interaction of a bioactive form of the JA hormone, jasmonoyl-*L*-isoleucine (JA-lIe), with the receptor CORONATINE INSENSITIVE 1 (COI1). This interaction leads to degradation of JASMONATE-ZIM DOMAIN (JAZ) proteins, resulting in the release of MYC transcription factors (TFs) that activate JA responsive genes. Repression of the JA signaling pathway occurs through interaction of JAZ repressors with NOBEL INTERACTOR OF JAZ (NINJA), which recruits the co-repressor TOPLESS (TPL) (Howe et al., 2018). In Arabidopsis, two major branches of the JA signaling pathway have been described. The MYC branch, controlled by MYC-type TFs, is associated with wound response and defense against herbivorous insects. The ethylene response factor (ERF) branch is associated with resistance to nectrophic pathogens. This branch is regulated by members of the APETALA2/ETHYLENE RESPONSE FACTOR (AP2/ERF) family of TFs, like ERF1 and OCTADECANOID-RESPONSIVE ARABIDOPSIS59 (ORA59) and leads to the transcriptional upregulation of *PLANT DEFENSIN1.2* (*PDF1.2*). The ERF branch is co-regulated by JA and ET signaling (Lorenzo et al., 2003, McGrath et al., 2005, Dombrecht et al., 2007, Pré et al., 2008).

Pathogens have developed various strategies to manipulate defense hormone signaling to render plants more susceptible to infection. The hemibiotrophic bacterial pathogen *Pseudomonas syringae* (*P. syringae*) produces the toxin coronatine (COR), which mimics JA-Ile in both structure and function. COR binds to COI1 leading to activation of JA signaling and suppression of SA-mediated defense against *P. syringae* (Katsir et al., 2008, Zheng et al., 2012). Effector proteins can also modulate JA signaling. HopBB1 and HopX1 from *P. syringae* interfere with JAZ repressor activity, leading to transcriptional activation of JA responses and thus promoting bacterial proliferation (Yang et al., 2017, Gimenez-Ibanez et al., 2014). The hemibiotrophic Arabidopsis fungal pathogens *Fusarium oxysporum f.sp, conglutinans* and *F. oxysporum f.sp. matthioli* produce isoleucine- and leucine-conjugated jasmonate, respectively, and exhibit reduced virulence in the *coi1* mutant, indicating that JA signaling is involved in promoting *Fusarium* infection (Cole et al., 2014). However, SA accumulation may not be involved in restricting *Fusarium* susceptibility as mutant plants defective in JA perception and SA accumulation (*coi1/NahG*) showed the same reduced level of infection with *F. oxysporum* as the *coi1* single mutant (Thatcher et al., 2009).

The Arabidopsis co-repressor family TPL/TPRs are involved in several plant processes, including JA and auxin signaling (Pauwels et al., 2010, Szemenyei et al., 2008), defense responses (Zhu et al., 2010), and meristem maintenance (Kieffer et al., 2006). TPL/TPR proteins contain several conserved domains. The N-terminal portion contains LIS1 homology (LisH), C-terminal to LisH (CTLH), and CT11-RanBPM (CRA) domains. The C-terminal portion contains two WD40 domains (Martin-Arevalillo et al., 2017). TPL/TPRs can interact with transcriptional regulators via short repression domains (RDs) in their protein sequences (Causier et al., 2012a). Several RDs have been identified and include the EAR motif, defined by a consensus sequence of either LxLxL or DLNxxP (Kagale et al., 2010), and the non EAR motif R/KLFGV (Ikeda and Ohme-Takagi, 2009). Proteins with an LxLxL motif have been found to interact with the N-terminal part of TPL/TPR proteins (Pauwels et al., 2010, Szemenyei et al., 2008) while the R/KLFGV motif mediates interaction with the first WD40 domain of TPL (Collins et al., 2019). However, little is known about the TPL/TPR domains involved in interaction with the DLNxxP motif.

Here we demonstrate that the *U. maydis* effector Jsi1 possesses a DLNxxP motif which is responsible for interaction with the second WD40 domain of TPL/TPRs, leading to activation of JA/ET signaling. This reveals a novel fungal strategy to increase host susceptibility and suggests that the second WD40 domain of Topless is involved in regulating JA/ET signaling. In Arabidopsis, *jsi1* expression leads to the induction of genes related to the ERF branch of JA/ET signaling. Jsi1 could activate the ERF branch by interfering with the binding between endogenous DLNxxP motif containing ERF TFs and TPL/TPRs. Arabidopsis plants expressing Jsi1 for 2 days were more susceptible to infection with *P. syringae*, indicating that Jsi1-dependent induction of JA/ET signaling would lead to biotrophic susceptibility. The identification of unrelated effector proteins possessing a DLNxxP motif in different fungal species and validation of the interaction between *M. oryzae, S. scitamineum*, and *S. reilianum* effectors with TPL/TPRs indicate a converging strategy to manipulate this central signaling hub in plants.

## Results

### Jsi1 interacts with the C-terminal part of Topless via its EAR motif

We screened putative *U. maydis* effector proteins for the presence of the DLNxxP EAR motif and identified the gene *jsi1* (*UMAG_01236)*, located in cluster 2A (Fig. S1a) (Kämper et al., 2006). The EAR motif of Jsi1 starts at amino acid 39 and is adjacent to the signal peptide cleavage site. As is characteristic for an effector gene, *jsi1* is transcriptionally induced during biotrophy and its expression peaks four days post infection (dpi) (Table 1, Fig. S1b). To test if Jsi1 without signal peptide (Jsi1_27-641_), can interact with TPL, we cloned three *Z. mays* TPL orthologs (*ZmTpl1*, *Zm00001d028481*; Z*mTpl2*, *Zm00001d040279*; *ZmTpl3*, *Zm00001d024523*) (Fig. S2a and b). Jsi1_27-641_ interacts with all three ZmTPLs in a yeast two-hybrid (Y2H) assay (Fig. 1a, Fig. S3a). To identify which TPL domain interacts with Jsi1, we split ZmTPL1 into its N-terminal portion comprising the LisH, CTLH, and CRA domains (ZmTPL1^Nt^) and C-terminal portion containing the WD40 domains (ZmTPL^Ct^). We tested both truncated proteins in a Y2H and found that Jsi1_27-641_ interacts with ZmTPL^Ct^ (Fig. 1a). To identify which of the two WD40 repeats could be responsible for this interaction, we further divided ZmTPL^Ct^ into two fragments, each containing a single WD40 repeat (WD40-1 and WD40-2) and tested them for interaction by Y2H. Jsi1_27-641_ specifically interacted with WD40-2 (Fig. 1b). To interact with ZmTPL/TPR proteins inside the plant nucleus, Jsi1 would need to be secreted and translocated into the host plant cell. One challenge to demonstrate the effector-host protein interaction is the relatively low expression level of *jsi1*. We created a solopathogenic *U. maydis* strain SG200 expressing full-length Jsi1 tagged c-terminally with 3xHA under the control of the strong biotrophy-induced *cmu1* promoter (SG200P_cmu1_:jsi1-3xHA) which allowed us to detect Jsi1 by western blot (Fig. 1c). We immunoprecipitated Jsi1-3xHA from infected maize seedlings using anti-HA antibody beads and were able to specifically detect co-immunoprecipitated endogenous maize TPL/TPR proteins by western blot (Fig. 1c, Fig. S4). To test the specificity of the Jsi1 / maize TPL/TPR protein interaction *in planta*, we mutated the DLNxxP EAR-motif in Jsi1 to AHNxxP (Jsi1m). We found that Jsi1m was stably produced *in planta* but did neither interact with ZmTPL1 in Y2H nor *in planta* Co-IPs assays (Fig. 1a and c), indicating a critical role for the DLNxxP motif in the interaction between Jsi1 and TPL/TPR proteins.

**Table 1.**
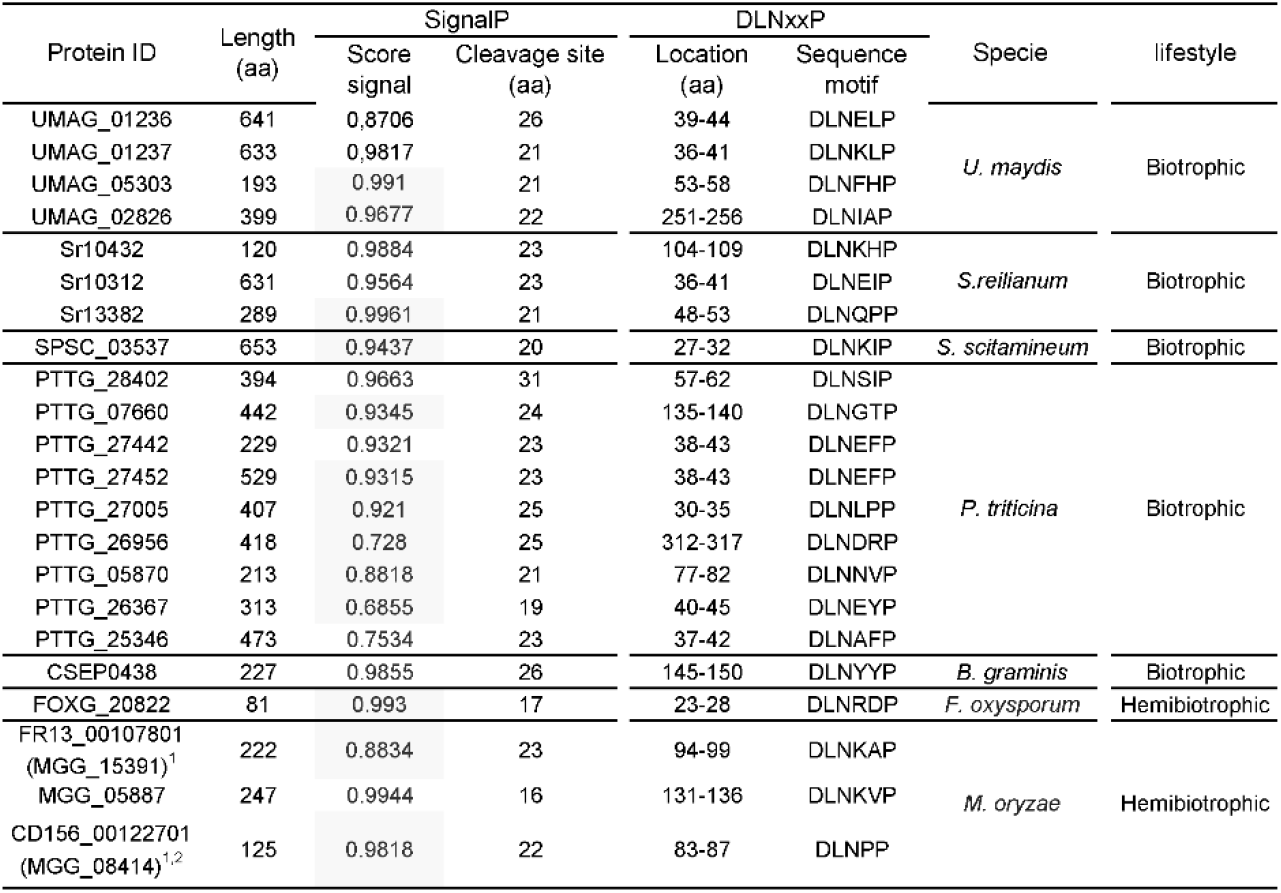
Fungal effector proteins possessing an DLNxxP motif. To identify putative secreted effector proteins with a DLNxxP motif, we performed a bioinformatic analysis where we selected proteins possessing the DLNxxP motif that are predicted to be secreted using SignalP-5.0 (http://www.cbs.dtu.dk/services/SignalP/) where the score signal value indicates the likelihood probability to get a secrete signal peptide, do not possess transmembrane domains using TMHMM-2.0 (http://www.cbs.dtu.dk/services/) and do not possess protein domains related with enzymatic activities using InterPro (https://www.ebi.ac.uk/interpro/beta/). 1 FR13_00107801 (*MGG_15391*) and CD156_00122701 (*MGG_08414*) were described in a comparative genomic project where different *M. oryzae* isolates where sequenced (Chiapello et al., 2015). ^2^ MGG_08414 was included in the analysis as it also possessed a DLNxxP-like motif to test if this motif can also interact with TPL/TPRs proteins.

**Figure 1.**
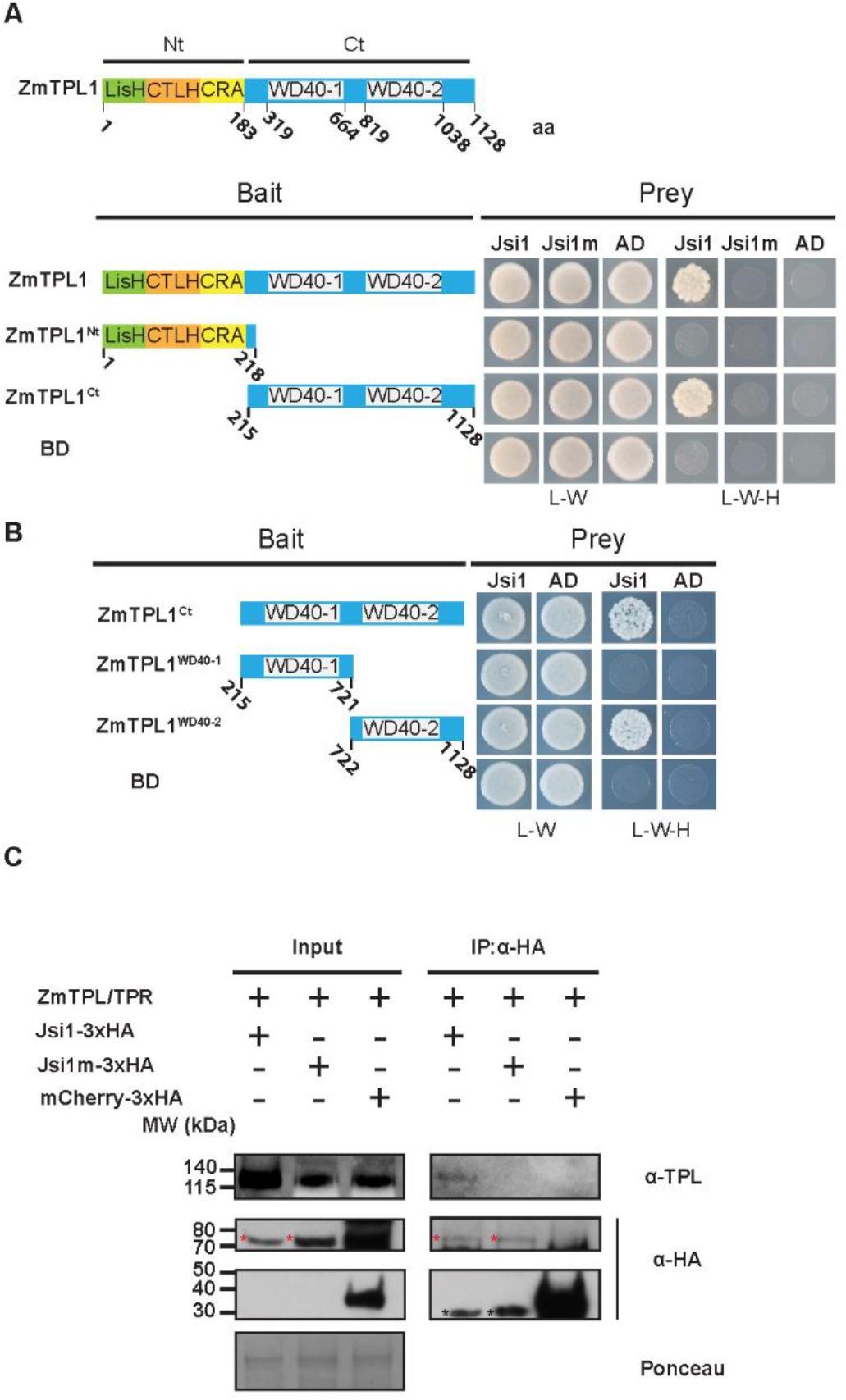
Jsi1 interacts with the second WD40 domain of ZmTPL1 through its DLNxxP motif (**A**) Y2H assay with Jsi1_27-641_ or the mutant version Jsi1m27-641 as prey and full-length ZmTPL1 or its N- and C-terminal regions (ZmTPL1^Nt^ and ZmTPL1^Ct^) as bait. (**B**) We further divided the ZmTPL1 C-terminal region in two parts, each containing one of the WD40 repeats, (ZmTPL1WD40-1 and ZmTPL1WD40-2) and used as baits to test which WD40 domain interacts with Jsi1 in Y2H assay. As negative control of interaction we used eYFP protein fused to the GAL4BD (BD) and GAL4AD (AD). The letters -L, -W and -H indicate medium lacking leucine, tryptophan and histidine, respectively. (**C**) Co-IP assay showing that Jsi1 interacts with ZmTPL/TPRs *in planta*. We infected maize seedling with each of the *U. maydis* strains expressing Jsi1-3xHA, the mutated version of Jsi1 (Jsi1m3xHA) and mCherry tagged with 3xHA and performed a Co-IP using anti-HA antibody to pull-down the recombinant proteins. TPL-specific antibody shows that endogenous maize TPL/TPR proteins are co-purified with Jsi1-3xHA but not with Jsi1m-3xHA nor mCherry-3xHA. Red asterisks indicate the full-length proteins of Jsi1-3xHA while black asterisks indicate unspecific band in the Jsi1-3xHA and Jsi1m-3xHA lanes. Ponceau staining was used as loading control.

### Jsi1 is a secreted effector located in the nucleus of maize cell leaves

Secreted fungal effectors can be bioinformatically predicted based on the presence of an N-terminal secretion signal that allows for secretion via the endoplasmic reticulum–Golgi apparatus route. After secretion, these translocated effectors must pass through the biotrophic interphase before being translocated into the plant symplast (Lo Presti et al., 2015). To test whether Jsi1 is secreted, as bioinformatically predicted by SignalIP-5.0 (http://www.cbs.dtu.dk/services/SignalP/), we ectopically integrated a 3xHA tagged version of Jsi1 under the control of the constitutive *otef* promoter into *U. maydis* strain AB33 (Brachmann et al., 2001). AB33 mimics pathogenic development in axenic culture and is therefore useful for studying effector secretion. As expected for a secreted protein, we were able to detect Jsi1-3xHA in the culture supernatant by western blot. Actin, which served as a lysis control, was only present in whole cell extracts (Fig. 2a). To confirm that Jsi1 is secreted *in planta*, we expressed a Jsi1-mCherry fusion protein in the strain SG200Δjsi1, where the endogenous *jsi1* locus is deleted. To increase protein levels for visualization, *jsi1-mCherry* was expressed under the control of the strong, biotrophy-induced *cmu1* promoter. Jsi1_27-641_-mCherry, which lacks the predicted signal peptide, was used as a negative control. We observed Jsi1-mCherry fluorescence around and outside the fungal hyphae while Jsi1_27-641_-mCherry was localized inside the fungal hyphae, indicating that Jsi1 is secreted by *U. maydis in planta* (Fig. 2b). Large tags, including mCherry and GFP, have been previously demonstrated to inhibit translocation of *U. maydis* effectors (Tanaka et al., 2015). To determine the subcellular localization of Jsi1 in maize cells, we transiently co-transformed a Jsi1_27-641_-mCherry construct driven by the 35S promoter together with a GFP-NLS construct as a nuclear marker into maize leaves. Using confocal microscopy one day after biolistic transformation, we found Jsi1_27-641_-mCherry signal inside the plant nucleus (Fig. 2c). To test whether ZmTPL1 colocalizes with Jsi1 in maize cells, we co-transformed a ZmTPL1-GFP expression construct with the Jsi1_27-641_-mCherry construct. ZmTPL1-GFP signal emission fully overlapped with the Jsi1_27-641_-mCherry signal in the nucleus, indicating the co-localization of both proteins (Fig. 2d).

**Figure 2.**
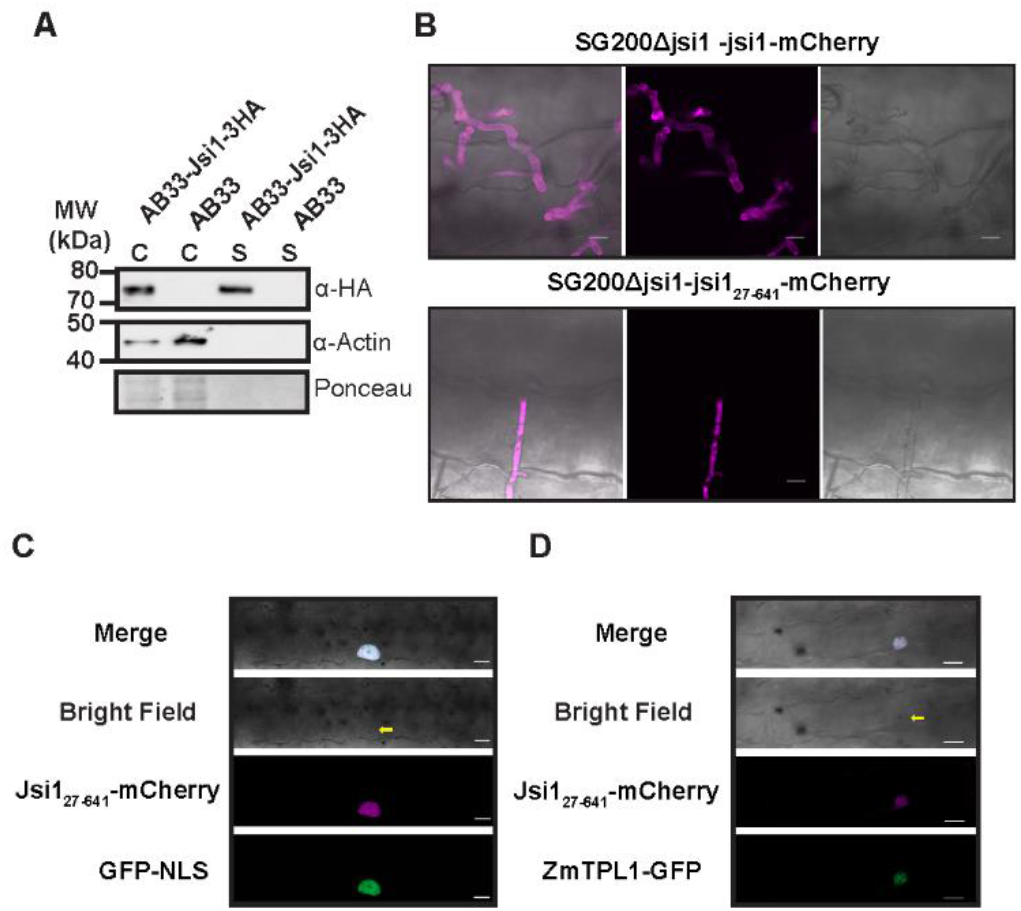
Jsi1 is a secreted effector that is targeted to the plant cell nucleus. (**A**) Jsi1 is secreted in axenic culture. We expressed Jsi1-3xHA fused protein constitutively in the *U. maydis* strain AB33. We extracted proteins from filamentous cells and culture supernatants and proteins were subjected to western blot analysis using anti-HA and anti-actin antibodies. We used actin as control of cell lysis, and it was only detected in whole cell extracts (“C”). Jsi1-3xHA was detected in whole cell extracts (“C”) and culture supernatants (“S”). (**B**) Jsi1-mCherry is secreted by *U. maydis in planta*. Confocal images of infected maize leaves 3 days post infection with SG200Δjsi1-Jsi1-mCherry and SG200Δjsi1-Jsi1_27-641_-mCherry (a non-secreted version of Jsi1). We observed mCherry fluorescence around and outside the fungal hyphae for the secreted version while we observed mCherry fluorescence from the Jsi1 version lacking the signal peptide solely inside the fungal hyphae. Scale bars, 10 μm. (**C**) Jsi1 localizes to the nucleus of maize cells. Maize cells expressing Jsi1_27-641_-mCherry and GFP-NLS as a nuclear marker after biolistic transformation of leaves. Confocal images show co-localization of the Jsi1-mCherry and GFP-NLS signals in the nucleus of the plant cell. (**D**) Jsi1 and ZmTPL1 co-localize in the nucleus of maize leaves cells. Maize cells expressing Jsi1_27-641_-mCherry and ZmTPL1-GFP. Confocal images show co-localization of both proteins in the nucleus. The yellow arrow indicates the transformed cell with the gold particle inside the nucleus. Scale bar, 20 μm.

*Jsi1* belongs to the *U. maydis* cluster 2a which was previously shown to cause a mild hypervirulence phenotype in maize when deleted (Kämper et al., 2006). To test whether Jsi1 contributes to virulence, we used the SG200Δjsi1 strain to infect maize seedlings. Plants infected with the mutant strain did not show any significant changes in symptom development in comparison to plants infected with the progenitor strain (Fig. S1c and d).

### Jsi1 activates JA/Ethylene and SA signaling in Arabidopsis

Topless is a highly conserved co-repressor in land plants (Causier et al., 2012b). Since Jsi1_27-641_ also binds to TPL and TPR proteins from *Arabidopsis thaliana* (Fig. S5a), we were able to study which pathways are manipulated by Jsi1 *in planta*. We generated two independent transgenic Arabidopsis lines expressing Jsi1_27-641_ fused to mCherry (mCh) under the control of the estradiol-inducible XVE system, XVE-Jsi1_27-641_-mCh 1 and 2, as well as an XVE-mCherry (XVE-mCh) control line. We confirmed expression of the transgenes after 6 hours of β-estradiol induction by western blotting (Fig. S5b) and then subjected induced samples to RNA-seq. PCA analysis of the resulting transcriptomes showed that replicates from the two Jsi1 lines group together and are separate from the replicates of the control line, which also clustered (Fig. S6a). We then calculated changes in gene expression in *jsi1* lines relative to the control line. Using cutoffs of fold change greater than 1.5 and a *P*-value < 0.05, we identified 1,090 differentially expressed genes (DEGs), 915 of which were upregulated and 175 were downregulated. The more than five times higher number of upregulated than downregulated genes is consistent with the model that Jsi1 interferes with the repressor-function of TPL/TPR proteins. The RNA-seq was validated by quantitative real-time PCR for a subset of up-regulated genes (Fig. S6b). Go-term analysis for biological processes show several categories related to “responses to different stimulus”. Within these, “responses to stress” (GO:0006950), “defense responses” (GO:0006952), “responses to external stimulus” (GO:0009605), and “response to biotic stimulus” (GO:0009607) were the major categories with 26, 15, 14, and 12 percent of the total DEGs, respectively. This indicates that Jsi1 induces plant immune responses in *A. thaliana* (Fig. S7a). In addition, we identified two Go categories related to hormone responses, “response to salicylic acid” (GO: 0009751) and “ethylene response genes” (GO:0009723).

In the ethylene response gene category, 26 out of 33 genes were upregulated and 14 of these were ERFs belonging to the AP2/ERF family of TFs. Seven of these 14 belong to the B3 group of the ERF subfamily, of which *ERF2*, *ERF5*, *ERF6*, and *DEWAX* (ERF107) have been associated with defense responses against necrotrophic infections and are positive regulators of the defense response gene *PDF1.2*, a well-known marker gene for the JA/ET signaling pathway (Ju et al., 2017, Moffat et al., 2012, McGrath et al., 2005) (Fig. 3a, Suppl. Table 1). Two other TFs of the B3 group, *ERF1* and *ORA59*, were found to be transcriptionally controlled by JA and ET and induce *PDF1.2* expression (Pré et al., 2008, Lorenzo et al., 2003). Even though we did not find *ERF1* and *ORA59* to be upregulated upon Jsi1 induction (Fig. S6b), a comparison of genes upregulated by Jsi1 and those found to be induced by *ERF1* and *ORA59* in previous studies show a 20 and 30 percent overlap, respectively (Fig. 3b). Go-term analysis of these overlapping genes show similar Go categories to DEGs from the Jsi1 expressing Arabidopsis lines where “response to stress”, “defense response”, “response to external stimulus”, and “response to biotic stimulus” are among the major categories with 57, 43, 41, and 41 percent of the total genes, respectively (Fig. S7b). Among the genes shared by Jsi1 with *ERF1* and/or *ORA59* we note the defense-related genes *OSM34*, *PR5*, and *PDF1.2* (Fig. S6b, Suppl. Table 1). In addition, we also found three ACC synthases (ACS) *ACS2*, *ACS6*, and *ACS11* and *MAP KINASE KINASE 9* (*MKK9*) which are involved in ethylene biosynthesis (Tsuchisaka et al., 2009, Xu et al., 2008), and *RGL3* a DELLA protein involved in the activation of JA-responsive genes (Wild and Achard, 2013) (Fig. 3a and Suppl. Table 1). Furthermore, Jsi1 induces the expression of 12 WRKY family TFs that were previously found to be induced by the necrotrophic fungal pathogen *Alternaria brassicicola* and/or by Methyl Jasmonate (MeJA) treatment or were found to be associated with necrotrophic resistance (Chen et al., 2010, McGrath et al., 2005) (Fig. 3a, Suppl. Table 1). Taken together, our results show that Jsi1 induces the expression of several ERFs from the ERF subfamily B3 group, genes related with ET synthesis, and defense genes including *PDF1.2*, indicating that Jsi1 induces the ERF branch of the JA/ET signaling pathways.

**Figure 3.**
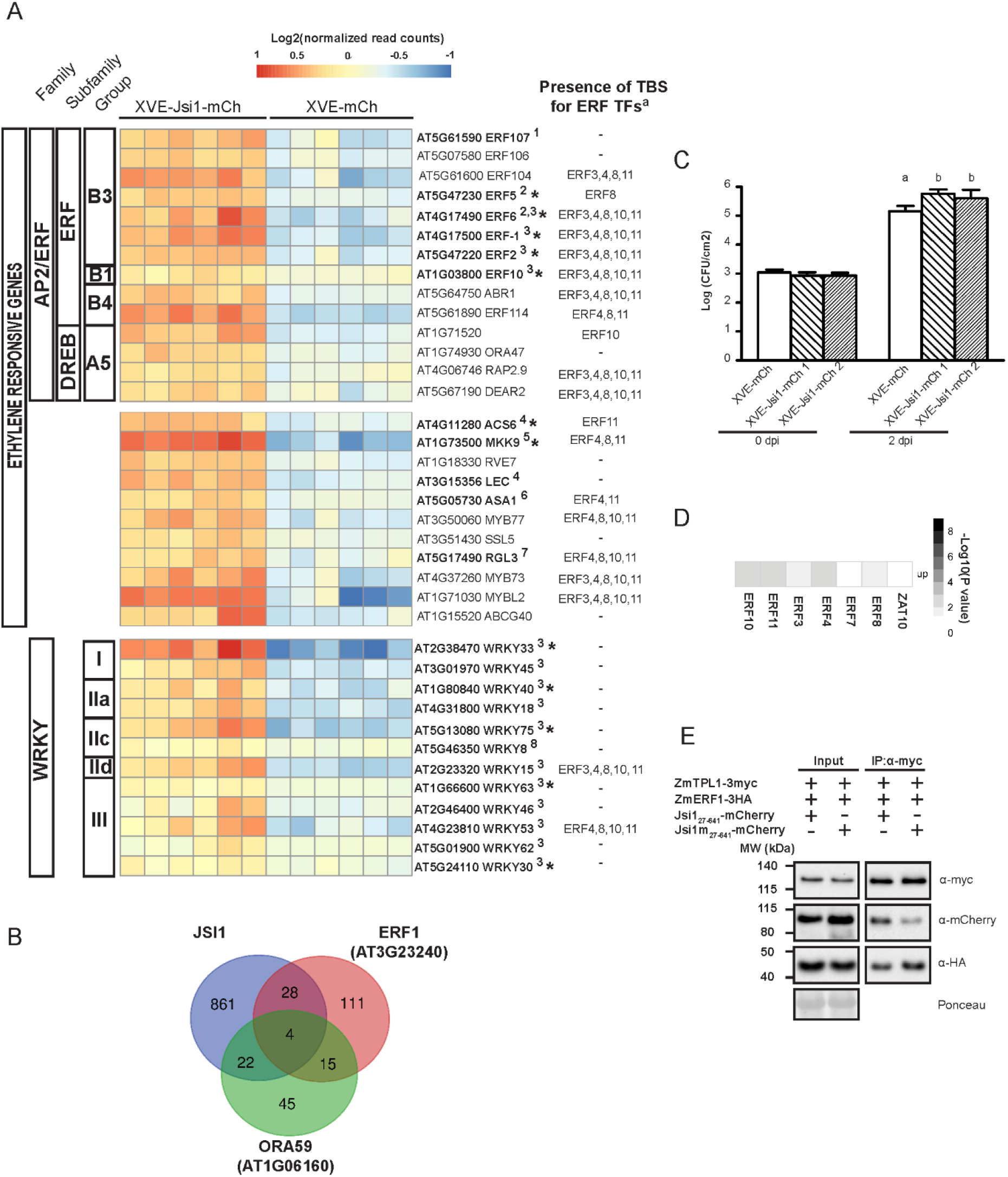
Jsi1 activates JA/ET signaling leading to biotrophic susceptibility. (**A**) Heat map from RNA-seq showing ethylene responsive and WRKY genes. Genes in bold and numbered in the heatmap were found associated with JA/ET signaling and/or response to necrotrophic resistance in the literature: ^1^ (Ju et al., 2017); ^2^ (Moffat et al., 2012); ^3^ (McGrath et al., 2005); ^4^ (Pré et al., 2008); ^5^ (Xu et al., 2008); ^6^ (Lorenzo et al., 2003); ^7^ (Wild and Achard, 2013); ^8^ (Chen et al., 2010). The asterisks indicate 12 out of 17 genes of which their expression levels were validated by qPCR. ^a^ Genes enriched in transcription binding sites (TBS) from ERFs with repression activity. (**B**) Venn diagram showing transcriptionally induced genes shared by Arabidopsis plants overexpressing either Jsi1, ERF1 or ORA59. (**C**) *Pst*. DC3000 proliferate better in Arabidopsis plants overexpressing Jsi1. We sprayed Arabidopsis lines with 150nM of β-estradiol 12 hours before *Pst* DC3000 inoculation. We collected infected leaves the same day of inoculation (0 day) and 2 days after inoculation to quantify bacterial proliferation. The graph shows one representative replicate of three repeated experiment. Statistical differences among the different genotypes were calculated by one-way ANOVA followed by Tukey, (*P* value < 0.05). (**D**) Genes upregulated upon Jsi1 induction are enriched in TBS of ERFs with a DLNxxP motif. Matrix summarizing the overlap enrichment between putative direct target genes of ERFs with repressor activity from previously available DAP-seq data and genes upregulated upon Jsi1 induction. Significance enrichment of TBS for each transcriptional regulator was determined by Fisher’s exact test (*P* value < 0.05). (**E**) Co-IP showing that Jsi1 can interfere with the ZmERF1 binding to ZmTPL1. We co-infiltrated Jsi1_27-641_-mCherry fusion protein with ZmERF1-3xHA and ZmTPL1-3xmyc. We performed Co-IP using anti-myc antibodies to pull-down ZmTPL1-3xmyc. WB analysis with anti-HA, anti-myc and anti-mCherry antibodies shows that Jsi1_27-641_ interferes with the binding between ZmERF1 and ZmTPL1 compared to Jsi1m_27-641_.

Regarding the Go category of SA response genes, 39 out of 40 genes were upregulated, including *GRX480*, *ALD1*, *WRKY70*, and *PR1* which have been described as SA-regulated genes (Herrera-Vásquez et al., 2014, Cecchini et al., 2015, Li et al., 2004, Durrant and Dong, 2004). In addition, we found two genes, *EPS1* and *PBS3*, involved in SA synthesis (Torrens-Spence et al., 2019) and regulators of the SA-induce immune response including *NPR3*, *NPR4*, and *NIMIN1* (Ding et al., 2018, Weigel et al., 2005) (Fig. S8). Activation of SA signaling could be due to recognition of the Jsi1-TPL interaction by the plant immune system as was previously reported for the *Fusarium oxysporum* effector six8 that interacts with TPL (Gawehns, 2014). Alternatively, the Jsi1-TPL interaction could interfere with the repressive activity of TPL-interacting transcriptional regulators involved in suppressing the SA signaling pathway.

### Jsi1 promotes biotrophic susceptibility in Arabidopsis

In order to asses whether the transcriptional changes observed in the JA/ET and SA signaling pathways correlated with changes in SA and JA hormone levels, we measured the levels of SA, JA, and JA-Ile, the bioactive form of jasmonate, in Arabidopsis shoots. The two Arabidopsis XVE-Jsi1-mCh lines expressing Jsi1 showed a significant increase in SA level compared to the XVE-mCh line while we could not detect JA-Ile or JA in either the XVE-Jsi1-mCh lines or in the control (Fig. S5c). The increased SA level could be a consequence of the upregulation of *EPS1* and *PBS3*, which are involved in SA synthesis (Torrens-Spence et al., 2019). On the other hand, the lack of JA and JA-Ile indicate that the activation of the ERF branch by Jsi1 is independent of the hormone. Activation of SA signaling should lead to repression of JA/ET signaling, as an extensive crosstalk between these two signaling pathways has been reported reviewed in (Caarls et al., 2015), and would increase resistance to biotrophic infection. To test if Jsi1 expression in Arabidopsis promotes biotrophic susceptibility, we tested the XVE-Jsi1-mCh and XVE-mCh control lines after estradiol-treatment for their susceptibility towards the hemi-biotrophic pathogen *Pseudomonas syringae* pv. *tomato (Pst*) DC3000. *Jsi1* expressing lines were more susceptible to *Pst* DC3000 infection than the control (Fig. 3c), indicating that activation of SA signaling in this context does not interfere with biotrophic susceptibility.

### Jsi1 can alter the repressing activity of ERF transcription factors

The activation of JA/ET and SA signaling by Jsi1 could be a consequence of its interaction with TPL/TPR proteins, leading to interference with the repressive activity of endogenous DLNxxP-containing transcriptional regulator. In *A. thaliana*, 67 transcriptional regulators were identified with a predicted DLNxxP motif including transcription factors from the B1 subfamily of ERF, C2H2, and JAZ families (Kagale et al., 2010). Fourteen of these were shown to directly interact with TPL/TPR proteins by Y2H assays, and the DLNxxP motif was essential for this interaction (Causier et al., 2012a). TFs which interact with TPL/TPRs are mainly negative regulators of transcription. We therefore focused on genes that are upregulated in the Jsi1 expressing lines that could be potential targets for DLNxxP-containing TFs. Using previously available DAP-seq data, we first determined genome-wide putative direct target genes of six ERFs from the B1 subfamily, ERF3, 4, 7, 8, 10, and 11, and ZAT10 from the C2H2 family (O’Malley et al., 2016). Next, we compared the list of putative target genes for each TF with those genes upregulated in the Jsi1 expressing lines and significant enrichment for each TF was evaluated by Fisher’s exact test (*P* value < 0.05). Except for ERF7 and ZAT10, transcriptional targets of the other ERFs were strongly enriched for genes derepressed by Jsi1. In total, 269 of the 915 genes upregulated by Jsi1 expression possess at least one transcription binding site (TBS) for an ERF (Fig. 3d and Suppl. Table 2). Go-term analysis of these up-regulated genes showed 26 categories, 24 of which were previously identified in the Go-term analysis of the Jsi1 overexpression RNA-seq data. Two categories of these were related with ethylene-activated signaling pathway (G0:0009873) and cellular response to ethylene stimulus (G0:0071369) whose genes were previously identified in the ethylene response genes category from the Jsi1 overexpression RNA-seq data (Fig. S7c). In fact, from the 26 upregulated genes identified in the RNAseq that respond to ethylene, 69% possess TBS for ERFs while from the 39 upregulated SA responsive genes only 33% possess TBS for ERFs (Fig. 3a and Fig. S8). This indicates that Jsi1 may regulate the expression of several ethylene responsive genes by altering the repressive activity of ERFs via interference with AtTPL/TPR proteins. Regarding to the SA responsive genes, some of them can be regulated by ERFs. However, the existence of others unknown transcriptional regulators whose interaction with TPL is altered by Jsi1 and possess a more relevant role in SA signaling activation cannot be excluded.

To test if Jsi1 can interfere with the interaction between ERFs and TPL/TPR proteins, we first verified that ERF3, 4, and 11 can interact with AtTPL in a Y2H assay. ERF3 possesses a DLNxxP motif, ERF11 possesses an LDLDLNFPP motif that combines two EAR motifs (DLNxxP and LxLxL), and ERF4 possesses the LxLxL and DLNxxP motifs in different regions of the protein. Both ERF3 and 11 interacted with the C-terminal portion of AtTPL while ERF4 interacted with both the N- and C-terminus (Fig. S9a) indicating that ERFs possessing a DLNxxP motif can interact with the C-terminal portion of AtTPL. We next cloned *ZmERF1* (*Zm00001d043205*) which possesses a DLNxxP motif and shares between 41% to 76% of protein sequence identity with AtERF3, 4, 8, 10, and 11. We found that ZmERF1 interacted with ZmTPL1 both by Co-IP, when transiently expressed in *N. benthamiana*, and in a Y2H, where the C-terminal portion of ZmTPL1 was required for interaction (Fig. S9b and c). In addition, we found that Jsi1_27-641_ interacted with ZmTPL1 in a Co-IP assay in *N. benthamiana* (Fig. S9d). To test whether Jsi1_27-641_ can interfere with the interaction between ZmERF1 and ZmTPL1, Co-IP experiments were performed between ZmERF1 and ZmTPL1 in the presence of either Jsi1_27-641_ or the version containing a mutated DLNxxP motif, Jsi1m27-641. We found that ZmERF1 is not as efficiently co-precipitated by ZmTPL1 in the presence of Jsi1 compared to the mutated version, indicating that Jsi1 weakens the interaction between ZmERF1 and ZmTPL1 in a DLNxxP motif-dependent manner (Fig. 3e). Taken together, our results suggest that Jsi1 interferes with the interaction between ZmERF1 and ZmTPL1 resulting in the expression of genes mainly connected with JA/ET signaling.

### Interaction with TPL via the DLNxxP motif is conserved in candidate effectors from different fungi

Effectors from Arabidopsis pathogens belonging to different kingdoms converge onto common host target proteins (Weßling et al., 2014, Mukhtar et al., 2011). As TPL/TPR proteins are highly conserved, we wondered whether effectors from various fungal pathogens with different host ranges had converged on a strategy to manipulate TPL/TPR signaling. We performed a motif search analysis across published proteomes of plant pathogenic fungi to identify putative secreted effector proteins with a DLNxxP motif. We searched for effectors with DLNxxP in the smut proteomes of *U. hordei*, *U. bromivora*, *S. scitamineum* and *S. reilianum* and identified 4 additional effector candidates (Table1). In addition, we performed the same search in other proteomes of plant pathogen fungi with different lifestyles and from different fungal divisions. Based on sequence availability, we selected the obligate biotroph *Puccinia triticina* (*P. triticina*) from the Basidiomycota division and the biotrophic *Blumeria graminis* (*B. graminis*), the hemibiotrophic *Fusarium oxysporum* (*F. oxysporum*) and *M. oryzae*, and the necrotrophic pathogens *Botrytis cinerea (B. cinerea*), *Sclerotinia sclerotiorum* (*S. sclerotiorum*), and *Bipolaris maydis* (*B. maydis*) from the Ascomycota division. In all of the pathogens evaluated, with the exception of the necrotrophic pathogens, we found at least one predicted secreted protein possessing a DLNxxP motif after the secretion signal (Table 1), indicating that effectors possessing a DLNxxP motif are mainly found in pathogens with biotrophic and hemibiotrophic lifestyles. To test if these effectors can interact with TPL, we selected *Sr10312* from *S. reilianum*, *SPSC_03537* from *S. scitamineum*, and FR13_00107801 (*MGG_15391*) from *M. oryzae*. In addition, we included a second *M. oryzae* effector, CD156_00122701 (*MGG_08414*) that contains a DLNxxP-like motif DLNxP to test if this motif can interact with TPL. *MGG_15391* and *MGG_08414* are expressed 24 hours after infection of rice, indicating that they may have a role during the biotrophic stage of *M. oryzae* (de Guillen et al., 2015, Shimizu et al., 2019). As the two *M. oryzae* effectors belong to strains able to infect rice (*Oryza sativa*) and *S. scitamineum* infects sugarcane (*Saccharum hybrid*), we cloned a rice TPL gene *OsTPL1* (*Os03g0254700*) and a sugarcane TPL gene *Sh_TPR3* (*Sh_231H24*) (Fig. S2). We fused the effectors lacking their secretion signals, MGG_1539124-222, MGG_0841423-125, Sr1031223-631, and SC_0353721-653, to mCherry and OsTPL1, ZmTPL1, and Sh_TPR3 to GFP. We performed Co-IP assays by co-expressing each effector with their respective host TPL in *N. benthamiana*. MGG_1539124-222, Sr1031223-631, and SPSC_0353721-653 co-immunoprecipitated with TPL proteins from rice, maize, and sugarcane, respectively (Fig. 4). Thus, our data indicate that TPL is a conserved target of effectors from hemibiotrophic and biotrophic fungal pathogens in the Basidiomycota and the Ascomycota divisions and mediated by the DLNxxP motif.

**Figure. 4.**
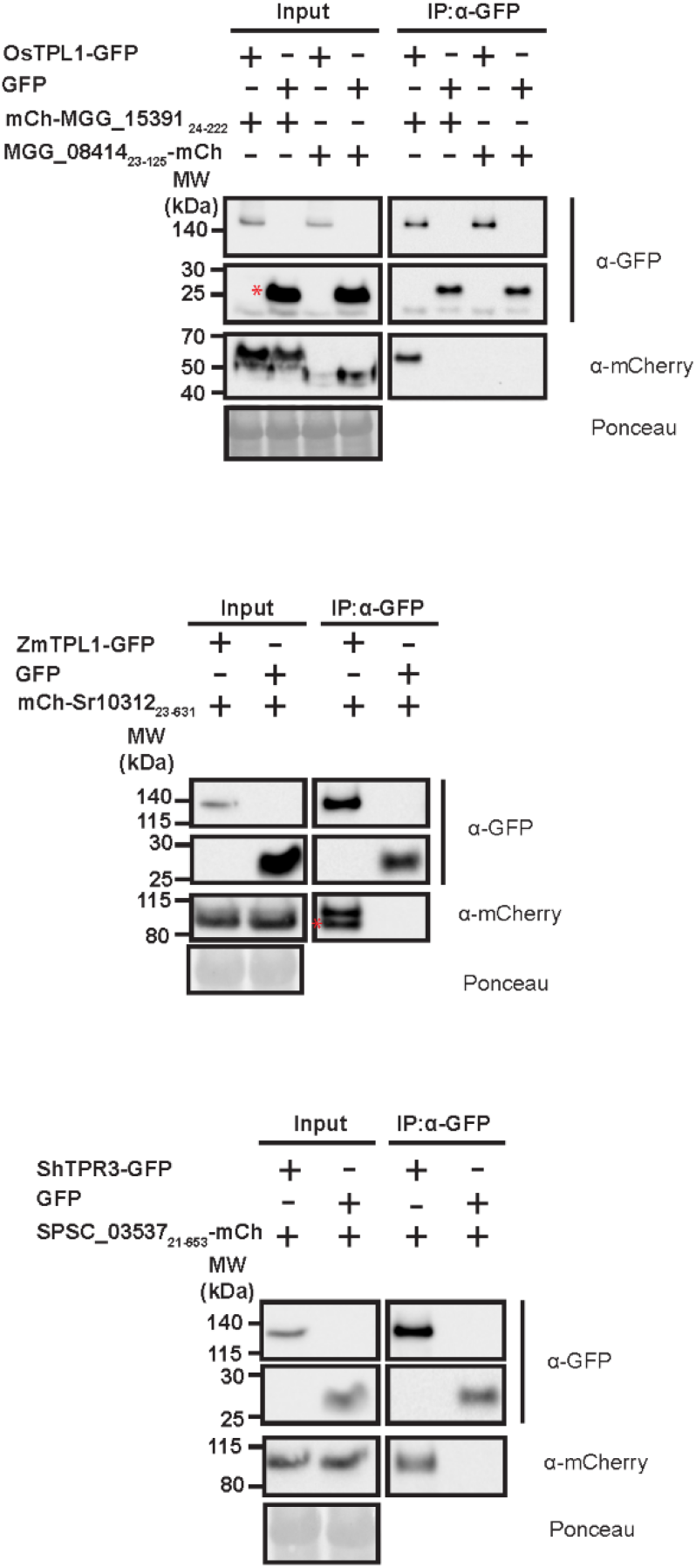
DLNxxP-motif containing effectors of different fungal pathogens are in complex with TPL. We fused mCherry (mCh) to fungal effectors and GFP to the TPL/TPR proteins. We co-infiltrated MGG_15391, MGG_08414, Sr10312 and SPSC_03537 with their respective TPL proteins in *N. benthamiana* leaves. We pulled-down TPL/TPRs-GFP with anti-GFP antibody and the co-immunoprecipitated proteins were detected with anti-GFP and anti-mCherry antibodies. GFP alone was used as negative control. The asterisks indicate the band belonging to GFP or Sr10312 full-length proteins respectively.

## Discussion

### Jsi1 hijacks the JA/ET signaling through interaction with the C-terminal part of TPL/TPR proteins

Pathogens have developed several strategies to activate JA-defense signaling, including the production of bioactive forms of JA or through effector proteins that can activate JA signaling (Howe et al., 2018). Infection of maize with *U. maydis* leads to the induction of JA biosynthesis and defense genes like defensins, hevein-like proteins, and chitinases, all of which attenuate the cell-death defense response. This suggests that induction of JA-defense signaling could be connected with repression of SA signaling (Doehlemann et al., 2008). However, how *U. maydis* induces JA-defense signaling remains unknown. Here, we identify a novel strategy in which *U. maydis* employs a DLNxxP-motif containing effector, *jsi1*, that activates the ERF branch of the JA/ET-defense signaling pathway by interacting with TPL/TPR proteins and which may contribute to biotrophic susceptibility. We show that interaction of Jsi1 with the C-terminal portion of different TPL/TPRs is dependent on the DLNxxP motif. In Arabidopsis, Jsi1 induces ERF TFs like *ERF2*, *ERF5*, *ERF6*, and *ERF107* which are associated with resistance to necrotrophic pathogens and the activation of the JA-defense signaling pathway (Ju et al., 2017, Moffat et al., 2012, McGrath et al., 2005). The induction of *PDF1.2* further supports the hypothesis that Jsi1 activates the ERF branch of the JA signaling pathway. In addition, we showed that *ERF1* and *ORA59* shared several upregulated genes with Jsi1 related to immunity. ERF TFs with a DLNxxP motif were described to interact with AtTPL/TPR (Causier et al., 2012a). AtERF3 and AtERF4 are active transcriptional repressors and the DLNxxP motif is essential for their activity (Ohta et al., 2001). In addition, ERF4 and ERF9 can suppress the expression of *PDF1.2* and are negative regulators of resistance to necrotrophic pathogens (Maruyama et al., 2013, McGrath et al., 2005). The significant enrichment of TBSs of several ERFs with repressor activity in 269 out of the 915 genes upregulated upon Jsi1 induction and the ability of Jsi1 to interfere with the interaction between ZmERF1 and ZmTPL1 indicate that ERFs with a DLNxxP motif are likely involved in repression of the ERF branch of the JA/ET signaling pathway. However, the existence of other unknown TFs with DNLxxP-like motifs that can also regulate JA/ET signaling cannot be excluded.

### The activation of SA signaling by Jsi1 expression does not lead to repression of JA/ET signaling in Arabidopsis

During the early stage of infection by *U. maydis*, plant immune responses connected with cell-death are activated. However, this immune response is attenuated 24 hours post infection, suggesting that *U. maydis* can inhibit programmed cell-death (Doehlemann et al., 2008). Jsi1 overexpression in Arabidopsis leads to activation of the SA signaling pathway as observed by the upregulation of several SA responsive genes like *GRX480*, *ALD1*, *WRKY70*, and *PR-1* (Herrera-Vásquez et al., 2014, Cecchini et al., 2015, Li et al., 2004, Durrant and Dong, 2004), upregulation of regulators of SA-mediated disease resistance such as *NPR3*, *NPR4*, and *NIMIN1* (Ding et al., 2018, Weigel et al., 2005), and an increase in total SA levels that may be connected with the transcriptional upregulation of two enzymes, *EPS1* and *PBS3*, involved in SA synthesis (Torrens-Spence et al., 2019). Activation of SA signaling could have several causes. Studying an effector function *in planta*, separated from the context of the rest of the effectome, may reveal complex responses derived from initial effector action and subsequent recognition responses by the plant immune system which would usually be counteracted by other effectors in the natural context. It was reported that interaction between six8, a *F. oxysporum* effector, with TPL leads to activation of SA-defense signaling in *A. thaliana* due to the Toll-like/interleukin-1 receptor-NB-LRR R protein SNC1, which guards TPL (Gawehns, 2014, Zhu et al., 2010). Therefore, the activity of Jsi1 on TPL may also trigger a guarding R gene. Another explanation is that the interaction between Jsi1 and TPL/TPR proteins could interfere with the repressive activity of unknown transcription factors with a DLNxxP motif, leading to activation of the SA signaling pathway. SA signaling based inhibition of JA/ET signaling has been previously demonstrated, reviewed in (Caarls et al., 2015) and ultimately would leads to increased resistance towards biotrophic interactions. However, Arabidopsis plants expressing Jsi1 are more susceptible to *Pst*. DC3000 infection. Furthermore, *PDF1.2* and several ERF TFs related with activation of JA/ET signaling are upregulated by Jsi1, indicating that JA/ET signaling cannot be repressed in this case by SA signaling. One explanation could be a potential role of EAR-motif containing ERFs in mediating the SA repression of the JA/ET signaling. As TPL is inhibited by Jsi1, these ERF TFs would lack their co-repressor and thereby be indirectly disarmed by Jsi1. It was previously reported that genes induced by MeJA that are antagonized by SA treatment are enriched in their promoter region for the GCC-box, the DNA binding motif of ERF TFs. In addition, it was shown that the GCC-box is sufficient for transcriptional suppression by SA and that SA leads to degradation of ORA59, a positive regulator of the ERF branch (Van der Does et al., 2013). Furthermore, the EAR-motif containing TFs ERF3, 10, and 11, can be transcriptionally induced by SA although single mutants of these ERFs do not show a JA/SA crosstalk phenotype, possibly due to redundancy (Caarls et al., 2017). In summary, Jsi1 activates both JA/ET and SA responsive genes but SA antagonism on JA/ET signaling, which is dependent on the ERFs – TPL/TPRs interaction, cannot be exerted as a consequence of the interaction of Jsi1 with TPL/TPRs (Fig. 5).

**Figure 5.**
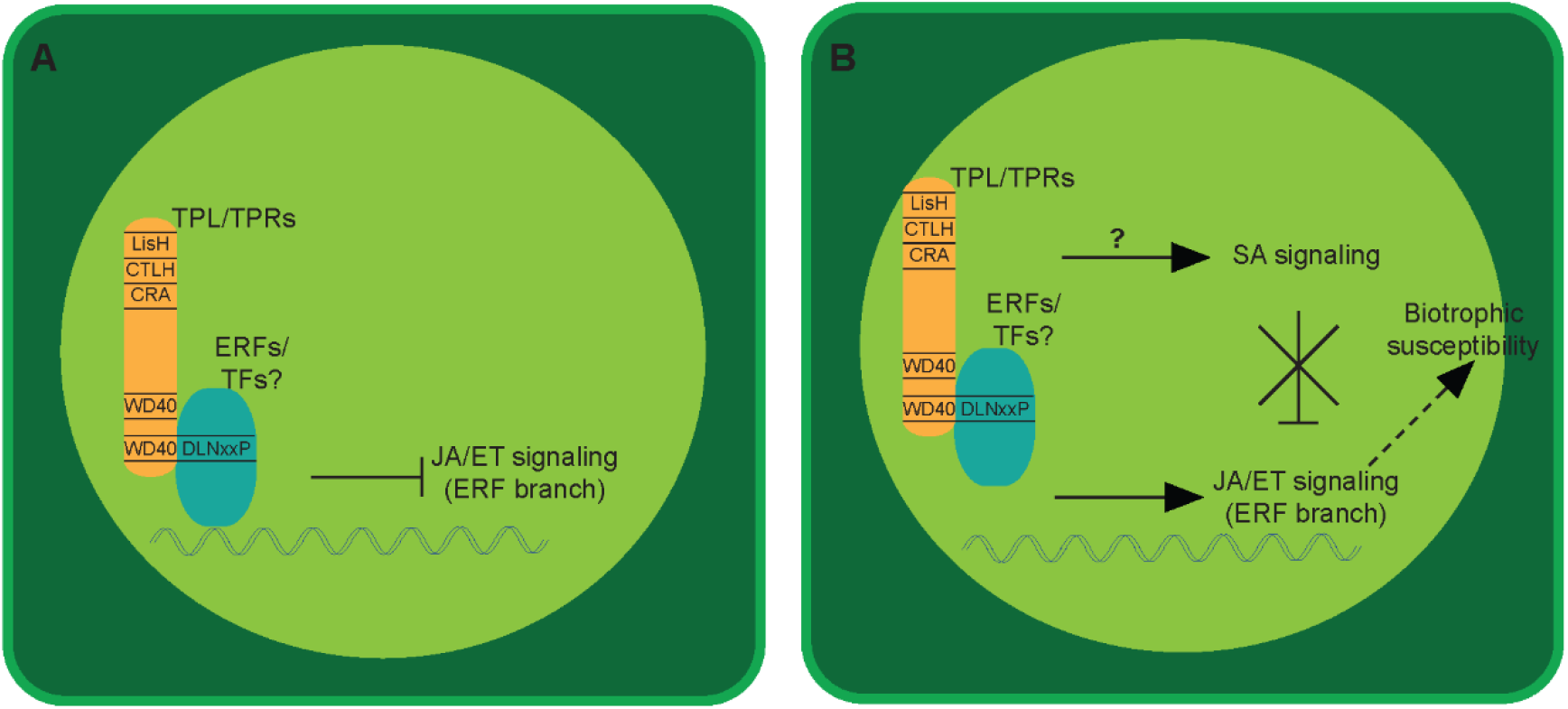
Jsi1 hijacks JA/ET signaling by interaction with TPL/TPRs. (**A**) In absence of fungal Jsi1 effector, interaction between plant ERFs and unknown TFs possessing a DLNxxP with the second WD40 domain of TPL/TPR corepressor proteins leads to repression of the ERF branch of the JA/ET signaling. (**B**) Jsi1 interferes with the binding between TPL/TPR proteins with ERFs and other unknown TFs leading to activation of the ERF branch of the JA/ET signaling. Activation of the SA signaling by Jsi1 could be due to activation of the plant immune system by recognition of the Jsi1-TPL/TPRs interaction by SCN1 as reported or another R protein that guards TPL/TPRs. On the other hand, SA signaling activation could be also due to Jsi1-dependent interference in the interaction between unknown TFs with TPL/TPRs. However, the SA signaling cannot repress the ERF branch of the JA signaling as repressive ERF TFs are blocked from binding to TPL/TPRs due to Jsi1 interference. Therefore, activation of the ERFs branch and a not fully deployed SA-defense pathway may lead to biotrophic susceptibility *in planta*.

### Arabidopsis, a model to study effector proteins with conserved host target/s

Most single effector deletions in *U. maydis* and many other fungal pathogens do not cause detectable impairments in virulence (Saitoh et al., 2012, Uhse et al., 2018). The possible reasons for this are manifold and could be due to technical limitations in measuring subtle differences in proliferation rates, artificial test conditions that do not reflect the selection pressures in nature, and functional redundancy with other effectors. One candidate effector that could play a redundant role to Jsi1 is *UMAG_01237*. It is located in the same gene cluster, possesses a DLNxxP motif, and interacts with ZmTPL1 in a Y2H assay (Fig. S3b). Nevertheless, the cluster 2a deletion mutant shows a weak hypervirulence phenotype. Additional effectors are therefore likely involved in manipulation of JA/ET signaling in maize (Kämper et al., 2006). Two other putative effector candidates, *UMAG_05303* and *UMAG_02826*, also possess a DLNxxP motif but did not show interaction with ZmTPL1 in a Y2H assay (Table 1, Fig. S3b). The identification of other functionally redundant effectors may prove difficult, as activation of the JA/ET signaling pathway might not be limited to manipulation of TPL/TPR activity or through the DLNxxP motif. Due to the limitations in studying effectors in their native systems, overexpression of effectors *in planta* is a commonly used approach. Working on a highly conserved host target like the TPL/TPR proteins allows for the use of the established model plant *A. thaliana* to study conserved mechanisms of hormone signaling, which we found to be exploited by Jsi1 to promote biotrophic susceptibility. Nevertheless, overexpression of effector proteins in a heterologous system can lead to effects that are not necessarily connected with the effector function. We observed that overexpression of Jsi1 in *N. benthamiana* leaves leads to cell death 5 days post infiltration and in Arabidopsis, depending on the β-estradiol concentration used to express Jsi1, cell-death is visible 3 days post β-estradiol treatment (Supp. Fig. S10). This could be due to SNC1 or other R proteins which may guard the Jsi1-TPL/TPRs interaction or through other activities of Jsi1. In the *in planta* secretion experiments, we used the strong cmu1 promoter instead of the native jsi1 promoter as native expression was too weak to enable detection of Jsi1-mCh by confocal microscopy. Therefore, fine-tuned expression of Jsi1 is required to induce JA/ET signaling and promote susceptibility without triggering cell-death.

### Effectors of diverse biotrophic and hemibiotrophic fungi convergently evolved

Plant host proteins targeted by effectors are under selective pressure to evade manipulation by the pathogen. On the other hand, if central regulators like TPL/TPRs interact with many endogenous host proteins via a specific motif, like the DLNxxP motif, it becomes nearly impossible to mutate the binding sites without tremendous fitness costs to the plant. This is likely why effectors from diverse biotrophic and hemibiotrophic fungi, including *Sr10312*, *SPSC_03537*, and *MGG_15391* from *S. reilianum*, *S. scitamineum*, and *M. oryzae*, respectively, may have convergently evolved the DLNxxP motif to specifically interfere with transcriptional control by TPL/TPRs. Jsi1 can serve as a prime example of effector biology, exploiting built-in signaling antagonisms in the host, targeting conserved signaling hubs, being possibly recognized in its action by the second layer of the plant immune system, and elucidating host biology by revealing a specific function of the C-terminal domain of TPL/TPR proteins in regulating JA/ET signaling in plants.

## Author Contributions

AD conceived the original research plan. AD and MD designed and coordinated the experimental work. DA, IF, JB, JM, KSC, KZ, MB, MD, LMSJ, RB, SU and YPH contribute to the experimental work. AD and MD wrote the manuscript.

## Conflict of Interest Statement

The authors declare that the research was conducted in the absence of any commercial or financial relationships that could be construed as a potential conflict of interest.

### Acknowledgments

We would like to acknowledge Dr. J. Matthew Watson for English correction on the manuscript. We would also like to acknowledge the GMI/IMBA/IMP service facilities, especially Dr. Robert Heinen and Ms. Zuzana Dzupinkova from the Molecular Biology services, Mr. Pawel Pasierbek from the BioOptics facility for his excellent technical support, and the Plant Sciences Facility at Vienna BioCenter Core Facilities GmbH (VBCF).

### Material and Methods

#### Plant material and growth conditions

*Zea mays* cv. Early Golden Bantam (EGB) (Olds Seeds, Madison, WI, USA) was used for infection with *U. maydis*. Maize was grown in a greenhouse (16 hr/8 hr light/dark cycle, 28°C/20°C). *Nicotiana benthamiana* (*N. benthamiana*) plants for CO-IP experiments were grown in a growth chamber (16 hr/8 hr light/dark cycle, 22°C, 60% humidity). Arabidopsis β-estradiol inducible lines XVE-Jsi1-mCh and control XVE-mCh lines expressing Jsi1_27-641_-mCherry-myc and mCherry-myc, respectively, were created by T-DNA insertion using agrobacterium infection in the Col-O DR5:GFP background. The estrogen receptor-based XVE system in the Arabidopsis lines is controlled by the CaMV 35S promoter and was cloned from the p1R4-p35S-XVE vector (Siligato et al., 2016). Arabidopsis plants were grown in a growth chamber (12 hr/12 hr light/dark cycle, 21°C, 60% humidity).

#### Strain and plasmids

All plasmids used in this work are provided in Suppl. Table 3. Plasmids by were generated by Golden Gate cloning as previously described (Lampropoulos et al., 2013).

*Ustilago maydis* strains used in this study were grown at 28°C in YEPSL medium (0.4% yeast extract, 0.4% peptone, 2% sucrose). SG200Δ*um01236 (jsi1)*, SG200, and AB33 strains were previously described in (Kämper et al., 2006, Uhse et al., 2018) and (Brachmann et al., 2001). All vectors and strains made in this study are available on request.

#### Secretion experiments in axenic culture and *in planta*

*U. maydis* strain AB33P_otef_ jsi1-3xHA was generated through insertion of plasmid pUG-P_otef_-Jsi1-3xHA into the *ip* locus of AB33 according to (Aichinger et al., 2003). We performed the secretion assay using strain AB33 according to (Brachmann et al., 2001). Briefly, filaments were induced by transferring to nitrate containing medium for 5 hr. We precipitated proteins in cell-free supernatant fraction with trichloroacetic acid (TCA)/Sodium deoxycholate (DOC) and precipitated proteins were subjected to western blot analysis. We used cell extracts and supernatants from an empty plasmid AB33 strain as a negative control to show specificity of the anti-HA antibody. We used actin as a control for cell lysis. The absence of actin in the supernatant fraction indicates that the presence of Jsi1-3xHA in the supernatant fraction was due to secretion and not cellular lysis. Mouse monoclonal anti-HA (Sigma Aldrich) and mouse monoclonal anti-actin (Invitrogen) antibodies for Western blot.

To visualize protein secretion *in planta*, we generated the SG200Δjsi1P_cmu1_Jsi1-mCherry strain by integrating *Jsi1*-mCherry under control of the cmu1 promoter in the *ip* locus using the integrative plasmid pUG. In addition, we built a non-secreted version of the Jsi1-mCherry strain by removing the predicted secreted signal (SG200Δjsi1P_cmu1_Jsi1_27-641_-mCherry). We independently infected both strains in 7 days old maize seedlings. 3 days post infection (dpi), mCherry fluorescence signal was detected using a Zeiss Axio observer Z1/7 confocal microscope.

#### Biolistic transformation of maize

We bombarded 6 day old maize leaves with 1.6 μm gold particles coated with Jsi1-mCherry and GFP-Nuclear Localization Signal (NLS) or Jsi1-mCherry and ZmTPL1-GFP using 5 μg of each plasmid as described by (Djamei et al., 2011). All proteins were expressed under the control of the CaMV35S promoter. Fluorescence emission was observed 1 day after transformation by confocal microscopy.

#### Virulence assay in maize

Three independent mutants in the *jsi1* locus (SG200Δ*jsi1-*1 to 3) were previously generated by (Uhse et al., 2018). To examine whether *jsi1* contributes to virulence, we assayed each of the mutant on 7 day old maize seedlings (100 plants per mutant). We quantified symptom development 12 dpi using the scale described by (Kämper et al., 2006). As a reference, we quantified disease symptom development in maize seedlings infected with the progenitor SG200 strain. We grew the deletion strains on filamentation inducing charcoal plates to test each strain for its ability to form filaments, a prerequisite for infection in the host plant. Significant differences between *jsi1* mutant strains and the SG200 progenitor were calculated using Fisher’s exact test. Multiple testing corrections were performed using the Benjamini-Hochberg algorithm. We repeated the experiment twice.

#### Arabidopsis *Pst* DC3000 infection assay

We performed *Pst* DC300 infection assay as described by (Liu et al., 2015). Briefly, we sprayed 4 week old Arabidopsis plants with a solution of 150 nM β-estradiol 12 hours before *Pst* DC3000 infiltration. We infiltrated leaves with a *Pst* DC3000 solution (OD_600_ = 0.0002) using a needleless syringe and inoculated plants were covered with a transparent lid to maintain humidity throughout the experiment. On the day of inoculation (0 dpi) and 2 dpi, leaf disks from six independent plants per genotype were collected using a 4 mm punch. We ground plant material in 2 ml Eppendorf tube containing 0.5 ml 10mM MgCl2 and 2 metal beads at 25 hz for 5 min. Serial dilutions were plated on King’s B agar medium and incubated at room temperature until colonies were visible. Bacterial proliferation was calculated as colony forming units (cfu) per cm^2^ of leaf and visualized on a Log scale (cfu/cm^2^). Statistical differences among the different genotypes were calculated using Tukey’s one-way ANOVA and *P* values < 0.05 were considered significant. The experiment was repeated three times.

#### Yeast two hybrid assay

We performed Y2H assays with the Mathmaker™ GAL4 Two hybrid system (Clonetech®) following the manufacturer’s protocol. We fused in frame to the GAL4 activation domain of the prey vector pGG446 (modified version of pGADT7) the genes *Jsi1*_*27-641*_, *Jsi1m*_*27-641*_, *UMAG_01237*_*22-633*_, *UMAG_05303*_*22-193*_, *UMAG_02826*_*23-399*_ (without signal peptide), *ERF3* (*AT1G50640*), *ERF4* (*AT3G15210*), *ERF11* (*AT1G28370*), and *ZmERF1 (Zm00001d043205*). We fused to the GAL4 binding domain from the bait vector pGG187 (modified version of pGBKT7) the genes *ZmTPL1* (*Zm00001d028481*), *ZmTPL2*, (*Zm00001d040279*), *ZmTPL3*, (*Zm00001d024523*)*, TPL* (*AT1G15750*), *TPR1* (*AT1G80490*), *TPR2* (*AT3G16830*), *TPR4* (AT3G15880) and N- and C-terminal portions of the different topless orthologs. We fused YFP to both GAL4 binding and activation domains and used them as negative control for interaction. We transformed the combinations of pGG446 and pGG187 vectors carrying the different genes in the yeast strains Y187 (MAT α) and AH109 (MAT a), respectively. After mating between bait and prey strains, we selected diploid yeast for growth on (SD)-Leu/-Trp, (SD)-Leu/-Trp/-His, and (SD)-Leu/-Trp/-His/-Ade plates at 28°C for 4 days. We repeated the experiments twice from independent mating events.

#### Co-immunoprecipitation (Co-IP) assay in *Nicotiana benthamiana*

We infiltrated four to five week old *N. benthamiana* leaves with *Agrobacterium tumefaciens* carrying different genes cloned into an expression vector as described in (Ma et al., 2012). Briefly, *A. tumefaciens* were grown overnight in LB medium containing corresponding antibiotics. We mixed cultures carrying the different genes and diluted them in agrobacterium re-suspension medium to a final OD600 of 0.3. We infiltrated six leaves (3 plants, 2 leaves from each plant). Leaves were collected 2 dpi, shock frozen in liquid nitrogen, and stored at −80°C. Tissue was ground using a Mixer Mill MM400 (Retsch) at 30 hz for 90 seconds. We resuspended 450 mg tissue powder in 2 ml cold extraction buffer (50 mM Hepes pH 7.5, 100 mM NaCl, 10% V/V Glycerol, 1 mM EDTA, 0,1% V/V Triton X-100, 2% PVPP, 1 mM DTT, 1 mM PMSF, and EDTA-Free Protease Inhibitor cocktail (Roche)). We sonicated samples for 5 min, 15s on/15s off, medium intensity using a BioRuptor^®^ (Diagenode). Lysates were centrifuged at maximum speed for 10 min at 4°C and the supernatant was transferred to a new tube. The centrifugation step was repeated three times. 100 μl of supernatant was mixed with 4xLDS buffer and the remaining supernatant was incubated with 30 μl of anti-myc or anti-GFP antibodies coupled to magnetic beads (μMACS™ MicroBeads, Miltenyi Biotech) for 1 h at 4°C. We loaded the supernatants with the magnetic beads onto magnetic columns (μ Column, Miltenyi Biotech) and the flow through was discarded. Columns were washed 4 times with 300 μl cold IP buffer (extraction buffer without 2% PVPP) and proteins were eluted using 2x LDS buffer heated to 95°C. We detected the immunoprecipitated proteins with anti-MYC (Sigma Aldrich), anti-HA (Sigma Aldrich), anti-mCherry (Abcam), or anti-GFP (Miltenyi Biotech) antibodies depending on the experiment. In the case of the Co-IP to test whether Jsi1 interferes with the binding between ZmTPL1 and ZmERF1, immunoprecipitated ZmTPL1 was quantified to load equal amounts of ZmTPL1 protein. Ponceau staining was used a loading control. All Co-IP experiments were repeated three times.

#### Co-IP assay in maize

*U. maydis* strains SG200P_cmu1_jsi1-3xHA, SG200P_cmu1_jsi1m-3xHA, and SG200P_cmu1_-SP_cmu1_(1-22)-mCherry-3xHA were generated by integrating plasmids pUG-P_cmu1_jsi1-3xHA, pUG-P_cmu1_jsi1m-3xHA, and pUG-P_cmu1_-SP_cmu1_(1-22)-mCherry-3xHA into the *ip* locus of SG200, respectively. We infected 7 day old seedlings with each strain (30 plants per strains). Infected tissue was collected 7 dpi, shock frozen in liquid nitrogen, and stored at −80°C. The Co-IP protocol was the same as for *N. benthamiana* above but using 3 g of infected tissue resuspended in 25 ml of extraction buffer and 150 μl of specific anti-HA antibodies coupled to magnetic beads (μMACS™ MicroBeads, Miltenyi Biotech). We detected immunoprecipitated proteins using anti-HA and anti-ZmTPL/TPR antibodies. Ponceau staining was used a loading control. We repeated the experiment three times.

The TPL-specific antibody was created using a small peptide, CNEQLSKYGDTKSAR, selected from conserved regions of the TPL/TPR proteins and synthesized by the Protein Chemistry core facility of GMI/IMBA/IMP. The polyclonal antibody was produced in rabbit by Eurogentec (Belgium).

#### Arabidopsis RNA-seq sample collection

Arabidopsis seeds from XVE-Jsi1-mCh-1/2 and XVE-mCh lines were sown on a nylon mesh with 100 μm pores (SEFAR^®^) placed over square plates containing 1% agar, ½ MS salts (Duchefa), 1% (w/v) sucrose, 0.05% (w/v) MES (pH 5.7 with NaOH). Arabidopsis plants were grown vertically in growth chamber (16/8 hr light/dark cycle, 21°C, light intensity 50 ɥmol/m^2^/sec) for 7 days. The nylon mesh with the Arabidopsis seedlings was transferred to square plates with same media containing 5 μM β-estradiol and grown vertically in the same growth chamber for 6 hr. Approximately 30 mg of seedlings were transferred to a 2 ml Eppendorf tube, shock frozen in liquid nitrogen, and stored at −80°C. Three independent replicates for each genotype were collected. Mock treatment was only performed for the control line to confirm that the concentration of β-estradiol used for the experiment did not itself alter gene-expression. Expression of Jsi1 and mCherry were evaluated after β-estradiol treatment by western blot using anti-mCherry antibody.

#### Illumina library preparation

We performed total RNA extraction of each replicate using the RNeasy Plant kit (QIAGEN). mRNA from each replicate was isolated via Poly(A) RNA Selection kit (Lexogen) using 1 μg of total RNA. We prepared libraries using the NEBNext^®^ Ultra™ II RNA library Prep Kit for Illumina and Index primers from NEBNext Multiplex Oligos for Illumina (New England BioLabs) following manufacturer’s instructions. Sequencing was performed on an Illumina Hiseq2500 machine with single-end 50-bp reads.

#### RNA-seq analysis

We removed adapter sequences and performed quality trimming using Trimmomatic (Bolger et al., 2014). Reads were mapped to the reference genome using STAR, version 2.7.0e, (Dobin et al., 2013) with the parameter --outFilterMismatchNoverLmax 0.05. We input the bam files to R 3.5.1 using the package Rsamtools. We obtained the genome annotation from Araport11 and gene models and read counts per gene were obtained with the packages Genomic Features and Genomic Alignments, respectively. We removed low-expressed genes and analyzed 28843 for differential expression using DESeq2, after performing regularized log transformation (Love et al., 2014). We compared the induced XVE-Jsi1-mCh lines with control XVE-mCh lines with and without β-estradiol treatment and kept genes with log fold change (FC) > 1.5 and adjusted *P* value < 0.05.

We performed Go-Term analysis for biological process using the ThaleMine tool from Araport11 (https://apps.araport.org/thalemine/) (Krishnakumar et al., 2014). Heatmaps were created using the package ‘pheatmap’ - R (https://cran.r-project.org/web/packages/pheatmap/pheatmap.pdf) and venn diagram using the tool from (http://bioinformatics.psb.ugent.be/webtools/Venn/).

To assess the significance of enrichment for transcription factor binding sites, we first determined the direct target genes of *ERF3*, *4*, *7*, *8*, *10*, *11*, and *ZAT10* using previously published DAP-seq data (O’Malley et al., 2016). We overlapped each list of putative direct target genes with genes upregulated upon Jsi1 induction (FC > 1.5, *P* value <0.05). We determined the significance of the overlapping with a Fisher’s exact test using the R package GeneOverlap function newGOM (https://github.com/shenlab-sinai/GeneOverlap).

#### RT-PCR for RNA-seq validation

Total RNA was extracted from three independent replicates from each Arabidopsis line (XVE-Jsi1-mCh-1 and 2 and XVE-mCh) using the same protocol for RNA-seq samples. cDNA was generated from 1 μg total RNA from each replicate using the iScript cDNA synthesis kit (Bio-Rad). We performed qRT-PCR with a Lightcycler 96 (Roche) using FastStart Universal SYBR Green Master mix (Roche) according to manufacturer’s instructions. The relative amount of amplicons in the samples were calculated with the 2^−ΔΔ*C*t^ method (Livak and Schmittgen, 2001) with *actin2* (AT3G18780) serving as the reference gene according to (Czechowski et al., 2005). We calculated fold change in the expression level of each gene in the XVE-Jsi1-mCh lines compared to the XVE-mCh line and data is represented as the mean of six replicates (three from each Jsi1-mCh line). We calculated statistically significant differences in gene expression between the Jsi1-mCh lines and mCherry line using the Mann-Whitney test with *P* value <0.05. Primers used for RT-PCR are described in Suppl. Table 4.

We designed specific primer pairs for each gene with Primer-BLAST (https://www.ncbi.nlm.nih.gov/tools/primer-blast/) with the following criteria: product size between 70 and 250 bp, primer length between 17 and 25 nucleotides, melting temperature between 57 and 63°C and,when it was possible, primer pairs must be separated by at least one intron. Specificity of the primers was confirmed by the presence of a single peak in the melting curve. Average amplification efficiencies of each primer pairs were derived from the formula of *E* =(10(^−1/slope^)−1)*100 and shown in percentage as described by (Svec et al., 2015).

#### Phytohormone measurements

We grew Arabidopsis lines XVE-Jsi1-mCh 1 and 2 and XVE-mCh control lines over a nylon mesh in square plates for 7 days as previously described. Plants were sprayed with a 5 μM β-estradiol solution and incubated for six hours. Shoots were cut, immediately frozen in liquid nitrogen, and phytohormone extraction was performed as described by (Kusch et al., 2019). Briefly, we performed reversed phase separation of constituents using an ACQUITY UPLC® system (Waters Corp., Milford, MA, USA) equipped with an ACQUITY UPLC® HSS T3 column (100 mm × 1 mm, 1.8 μm; Waters Corp., Milford, MA, USA) followed by Nanoelectrospray (nanoESI) analysis. Phytohormones were ionized in a negative mode and identified in a scheduled multiple reaction monitoring mode with an AB Sciex 4000 QTRAP® tandem mass spectrometer (AB Sciex, Framingham, MA, USA). We performed mass transitions as described by (Iven et al., 2012) with modifications specified in Suppl. Table 5.

#### Identification of putative secreted effectors proteins with DLNxxP motif

We downloaded predicted proteins from *Puccinia triticina* 1-1 BBBD Race 1, *Botrytis cinerea* B05.10, *Fusarium oxysporum* f. sp. lycopersici strain 4287, *Magnaporthe oryzae* 70-15, *Blumeria graminis* f. sp. hordei DH14, *Sclerotinia sclerotiorum* 1980 UF-70, *Bipolaris maydis* c5, *Sporisorium reilianum* SRZ2, *Sporisorium scitamineum, Ustilago hordei* Uh4857-4, *Ustilago bromivora* UB2112, and *Ustilago maydis* 521 from EnsemblFungi (https://fungi.ensembl.org/index.html) or NCBI (https://www.ncbi.nlm.nih.gov/). To identified putative secreted effector proteins with a DLNxxP motif, we searched for the DLNxxP motif in all predicted proteins from the different fungi species using CLC Main Workbench-7.7.2 (QIAGEN). Then, we searched for proteins with a predicted secretion signal (SignalIP-5.0, http://www.cbs.dtu.dk/services/), no transmembrane domains (TMHMM-2.0, http://www.cbs.dtu.dk/services/). and no predicted enzymatic domains (InterPro, https://www.ebi.ac.uk/interpro/beta/).

---

**Figure S1.**
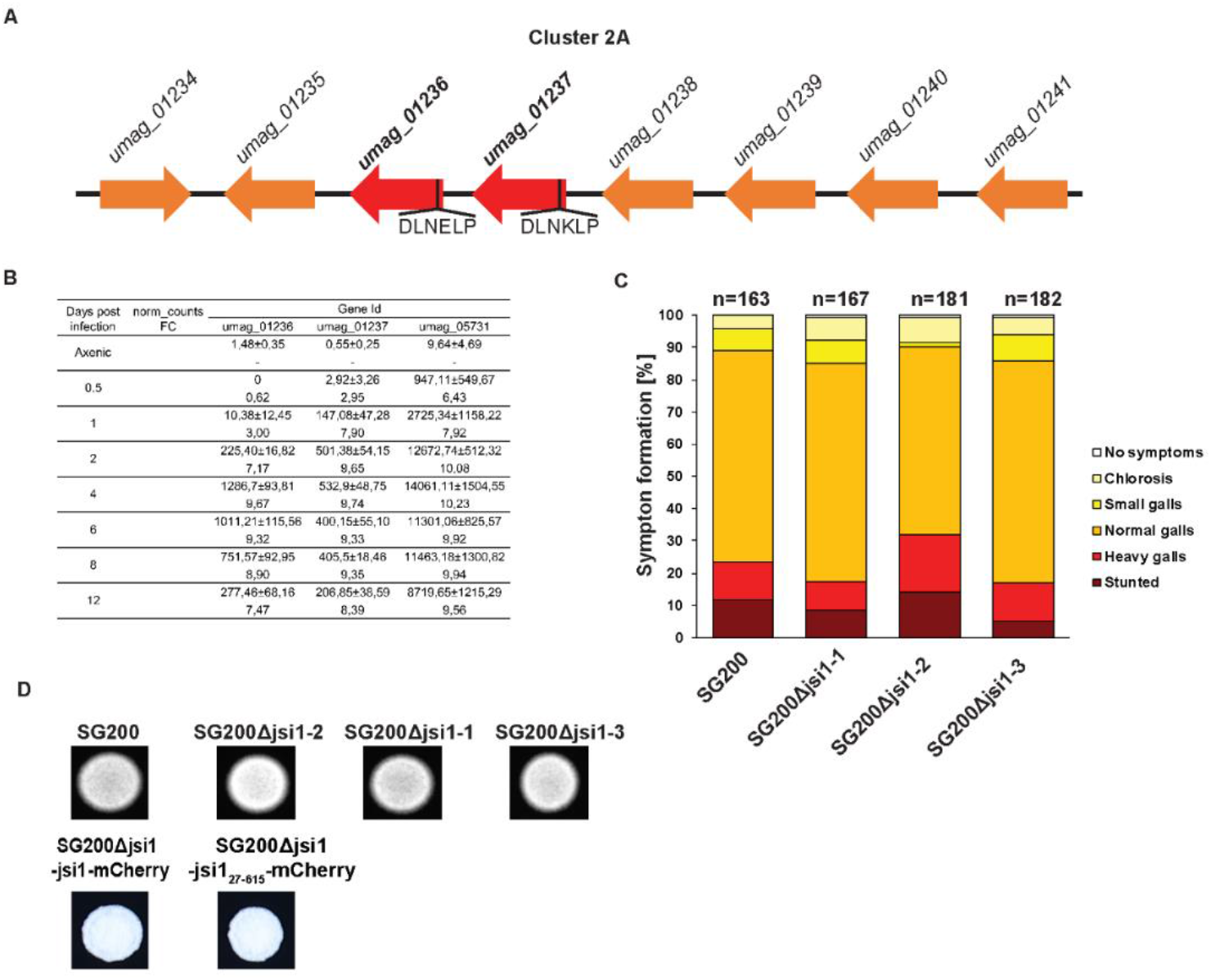
*jsi1* is part of effector cluster 2A and its deletion has no detectable contribution to the virulence of *U. maydis*. (**A**) Schematic representation of the *U. maydis* cluster 2a according to Kämper et al., (2006), *UMAG_01236* and *UMAG_01237* are in bold for being the only two genes of the cluster with a DLNxxP motif. (**B**) *UMAG_01236* and *UMAG_01237* are transcriptionally induced during infection. We extracted norm_count (normalized *U. maydis* reads counts) and FC (log2(Fold change)) from (Lanver et al., 2018). (**C**) Disease rating of Early Golden Bantam maize seedlings infected with *jsi1* mutants or progenitor strain SG200 respectively. We performed scoring of infected plants 7 days post infection. We compared the virulence phenotype of three independent mutants with the virulence phenotype of the progenitor strain. Numbers above the columns indicate total number of plants infected in two independent experiments. (**D**) We performed a filamentation test on charcoal containing plates for all the strains that we used for virulence analysis and for confocal imaging. Filamentation on charcoal plates indicates that the tested strains are not affected in their ability to form hyphae, a morphological prerequisite for virulence.

**Figure S2.**
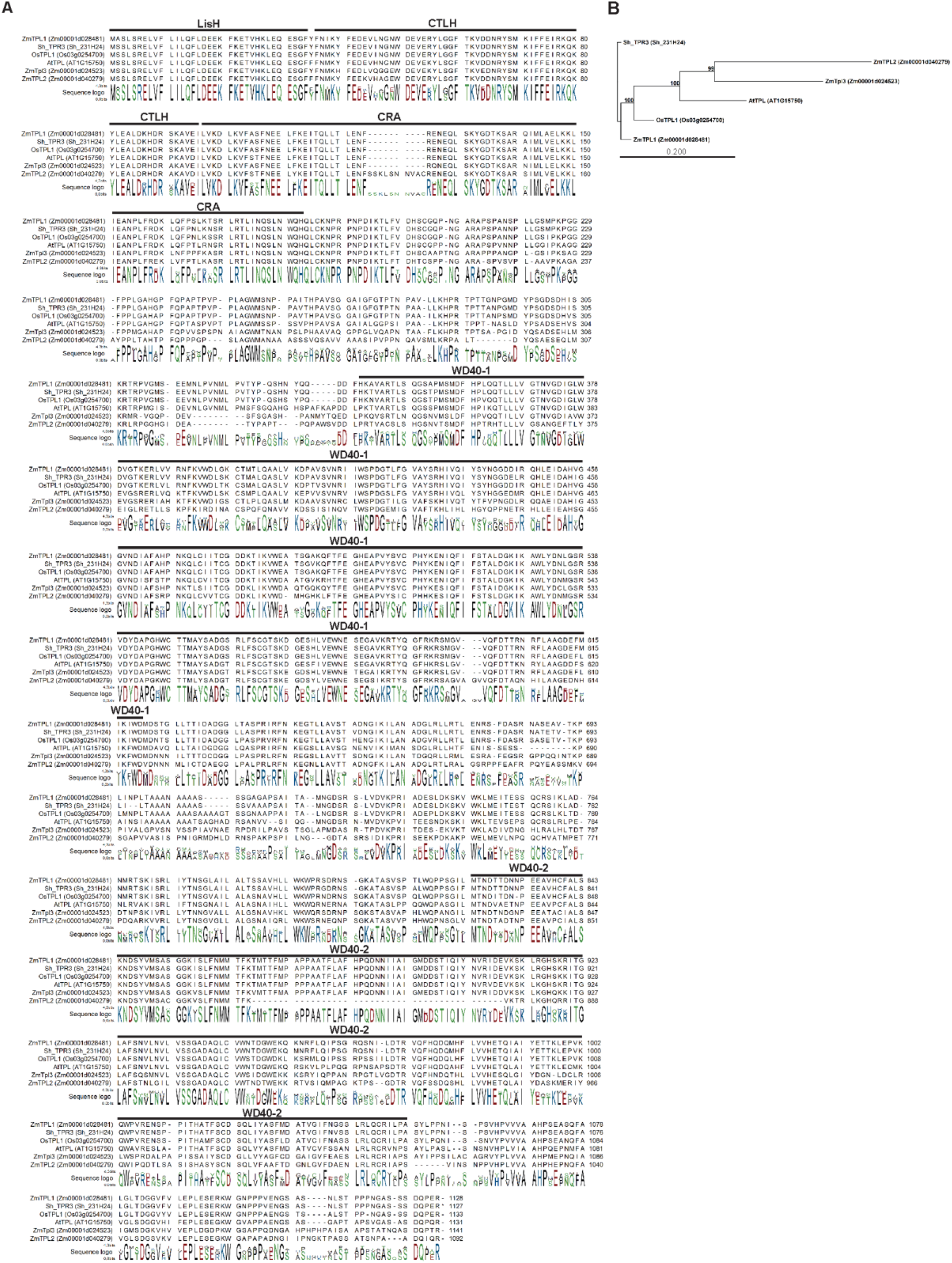
Phylogenetic analysis of TPL orthologs from different plant species. (**A**) Sequence alignment of TPL orthologs. Protein domains in ZmTPLs, OsTPL and ShTPR3 were defined according to the AtTPL domains identified in Martin-Arevalillo et al., (2017). (**B**) Phylogenetic tree of the TPL orthologs. Scale bar indicates Jukes-Cantor distance. Bootstrap values are shown in branches. At, *Arabidopsis thaliana*; Zm, *Zea mays*; Os, *Oryza sativa*; Sh, *Saccharum hybrid*. We generated sequence alignment and phylogenetic tree using CLC Main Workbench v 7.7.2.

**Figure S3.**
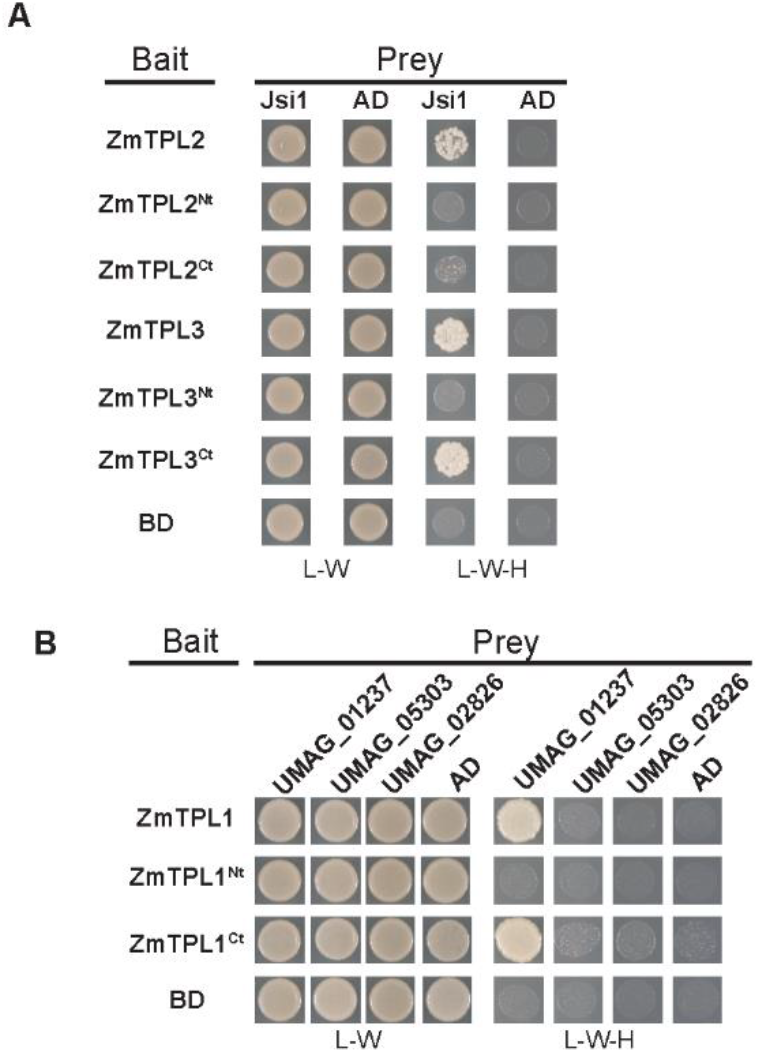
Jsi1 interacts with ZmTPL2 and ZmTPL3. (**A**) We used Jsi1_27-641_ as prey and ZmTPL2, ZmTPL3, and N and C terminal regions from both ZmTPLs (ZmTPL^Nt^ and ZmTPL^Ct^) as bait in a Y2H assay. (**B**) Umag_01237 interacts with the C terminal part of ZmTPL1 in Y2H. We used as prey the DNLxxP motif containing effectors Umag_01237, Umag_05303 and Umag_02826 without secretion signal. We used as bait the ZmTPL1 and its N and C terminal regions (ZmTPL1^Nt^ and ZmTPL^Ct^). We used eYFP protein fused to the GAL4BD (BD) and GAL4AD (AD) as negative control of interaction. The letters -L, -W and -H indicate medium lacking leucine, tryptophan and histidine, respectively.

**Figure S4.**
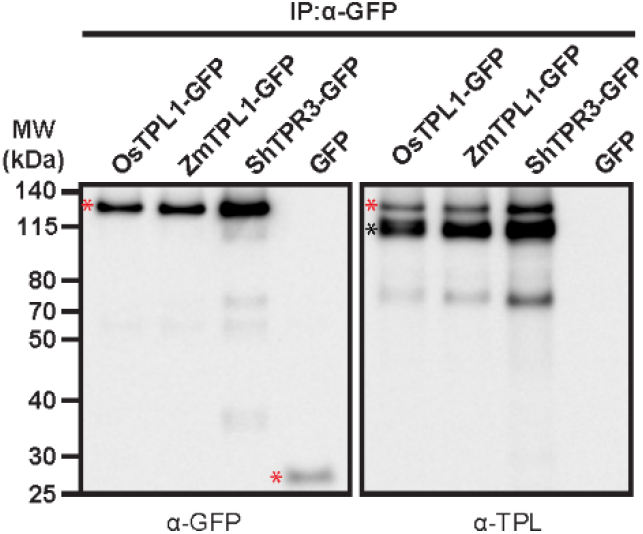
Topless-specific antibody can detect TPL/TPR proteins from different plant species. We fused ZmTPL1 (*Zea mays*), OsTPL1 (*Oryza sativa*) and ShTPR3 (*Saccharum hybrid*) to GFP and independently infiltrated in *N. benthamiana* leaves. We pulled-down the fused proteins from protein extracts using anti-GFP antibody. Western blot analysis with anti-GFP and TPL-specific antibody show that the TPL antibody can detect TPL proteins from different plants species. GFP was used as negative control to show specificity of TPL antibody by TPL/TPR proteins. The red asterisk indicates full-length of the different TPL/TPRs fused to GFP and GFP protein. The black asterisk indicates the TPL/TPR proteins from *N. benthamiana* that were pulled down by the TPL/TPRs fused to GFP as TPL/TPR proteins can form dimers and tetramers.

**Figure S5.**
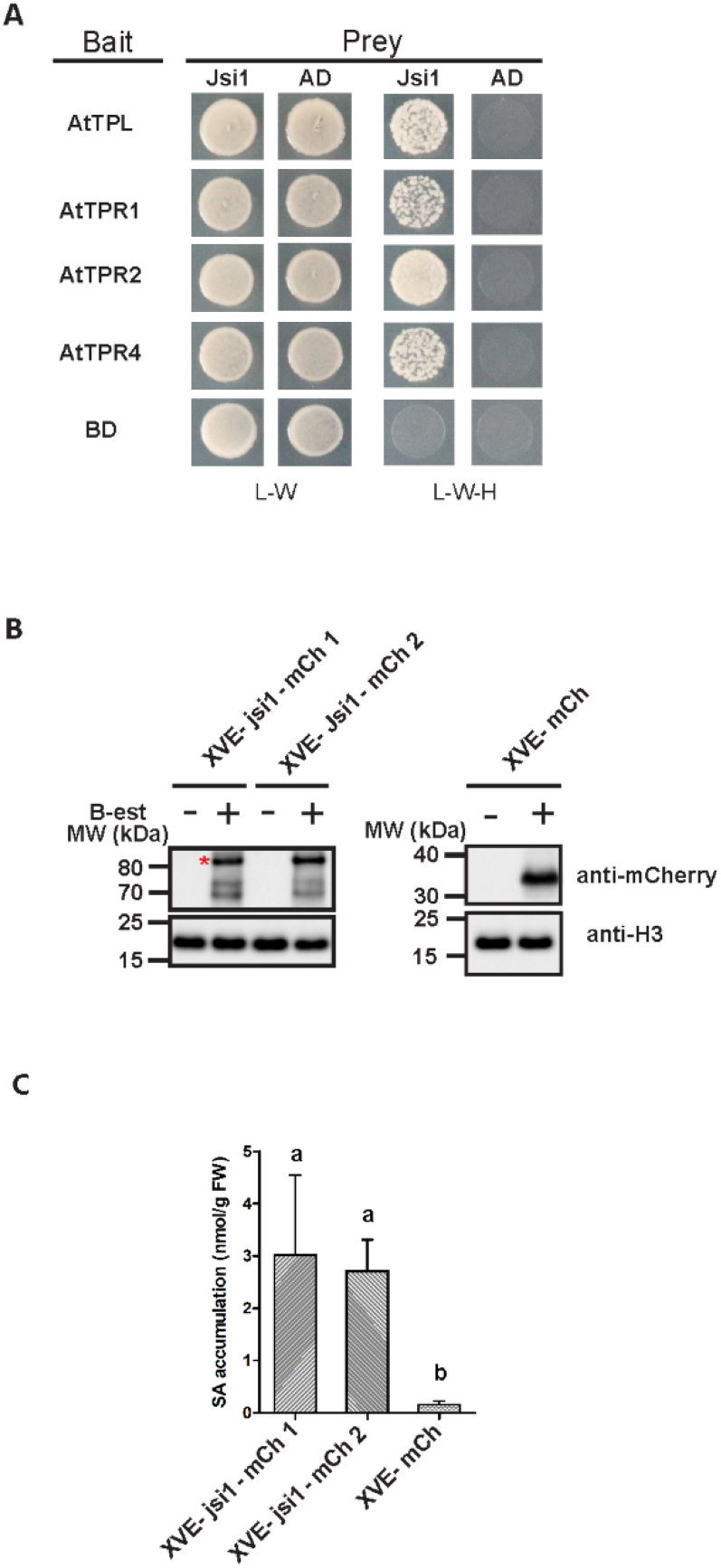
Jsi1 interacts with AtTPL/TPRs. (**A**) We used Jsi1_27-641_ as prey and AtTPL, AtTPR1, AtTPR2 and AtTPR3 as bait to test if Jsi1 can interact with the AtTPL/AtTPR proteins in Y2H assay. The eYFP protein was fused to the GAL4BD (BD) and GAL4AD (AD) to be used as negative control of interaction. The letters -L, -W and -H indicate medium lacking leucine, tryptophan and histidine, respectively. (**B**) Estradiol inducible XVE-Jsi1-mCh 1 and 2, and control XVE-mCh Arabidopsis lines. We treated Arabidopsis seedlings with 5μM of β-estradiol for 6 hours. We determined the expression levels of the proteins, Jsi1-mCherry or mCherry alone, using anti-mCherry antibody. Anti-Histone 3 antibody (Abcam) was used as loading control. Asterisk indicates full-length protein. (**C**) Total Salicylic acid content in shoots of fresh Arabidopsis seedlings. We sprayed XVE-Jsi1-mCh 1 and 2 and XVE-mCh lines with 5μM of β-estradiol to induce Jsi1-mCh and mCherry expression, respectively. After 6 hours post-induction we harvested shoots and subjected to LC-MS/MS for SA measurement. Each bar represents mean and standard deviation of five replicates. Significant differences between Arabidopsis XVE-Jsi1-mCh and XVE-mCh lines were calculated using Kruskal Wallis test followed by Dunn multiple comparison test (P < 0.05).

**Figure S6.**
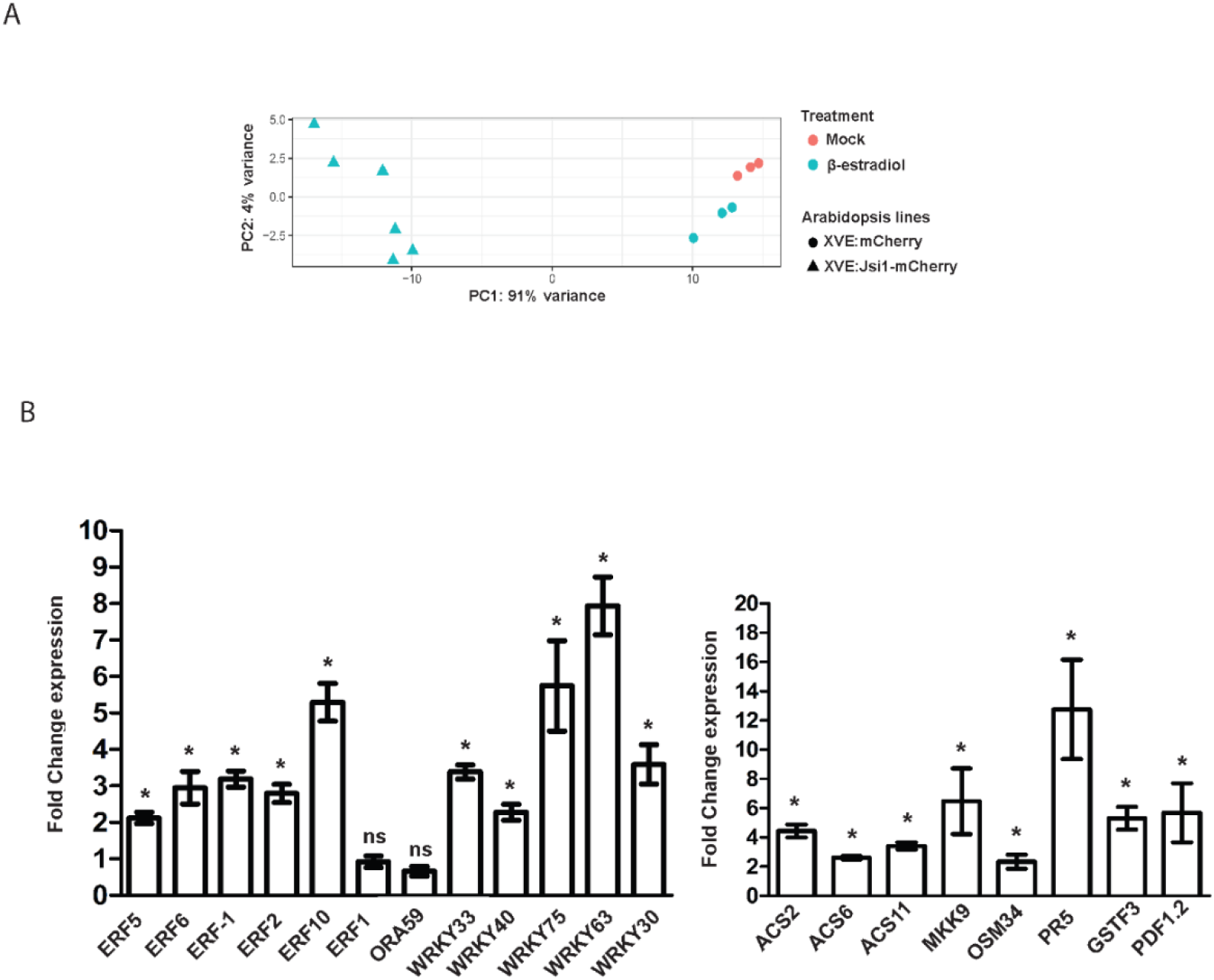
Principal component analysis (PCA) on RNAseq results and RT-PCR validation for some upregulated genes upon Jsi1 expression in *A. thaliana*. (**A**) The PCA demonstrates that RNA-seq libraries from induced XVE:Jsi1-mCh lines grouped together while libraries from XVE:mCh control line either treated with or without β-estradiol cluster together separately from the Jsi1 lines. (**B**) qPCR validation of genes from the RNA-seq. We selected upregulated ERFs, WRKYs and defense-related genes for q-PCR validation. *OSM34*, *PR5*, *GSTF3* and *PDF1.2* were found upregulated by *ERF1* and/or *ORA59*. Fold change is relative to the expression in the control mCherry line and normalized to the actin RNA expression values. Values shown are the means of six replicates (three from each XVE-Jsi1-mCh line), error bars indicate the SD. We evaluated statistically significant differences between gene expression in Jsi1 lines and the control mCherry line using Mann-witness test; *P* value <0.05. ns, not significant.

**Figure S7.**
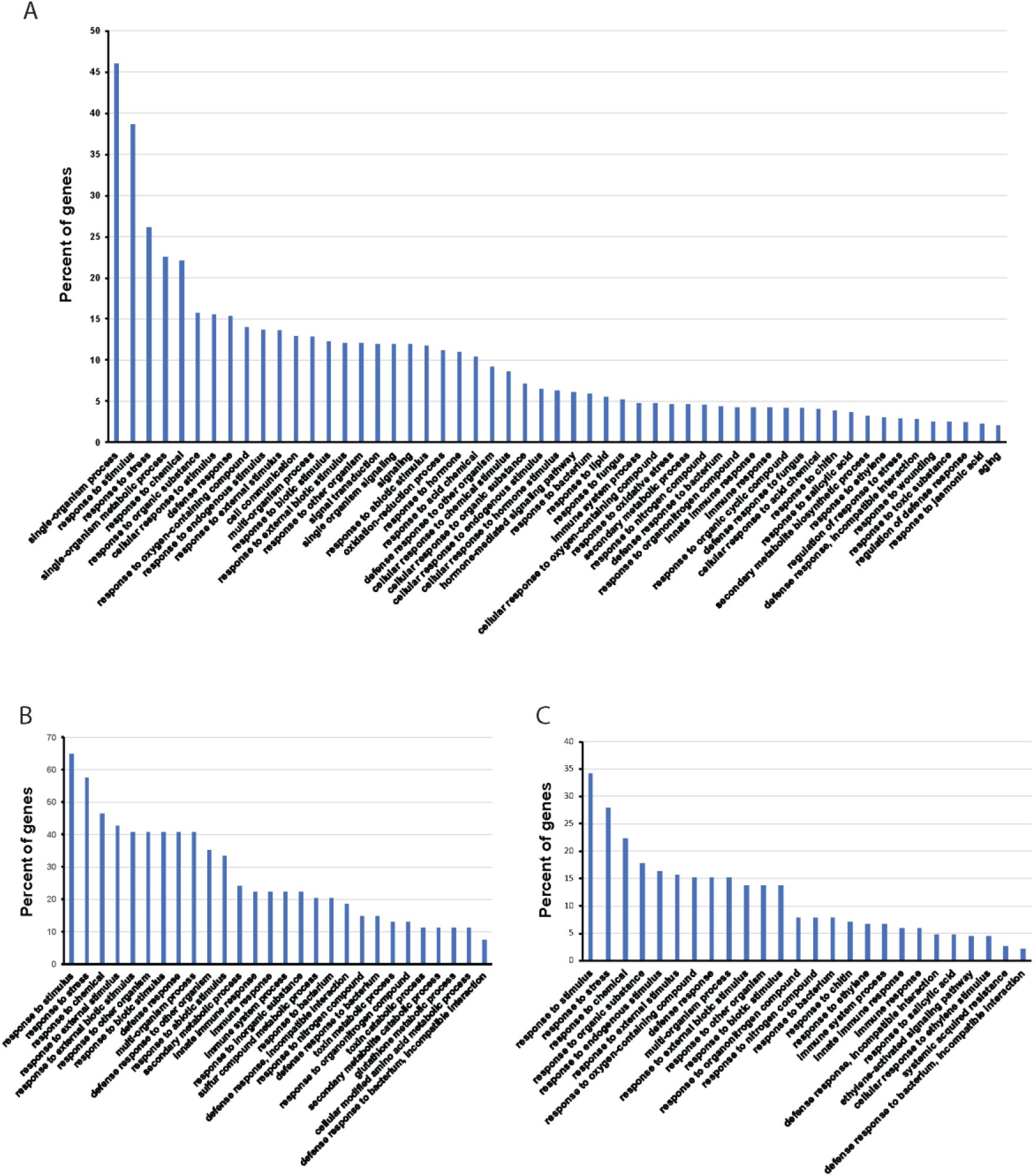
GO-term analysis for biological process of genes differentially expressed in Arabidopsis lines expressing Jsi1. (**A**) Go-term analysis for the 1,090 differentially expressed genes in lines expressing Jsi1. (**B**) GO analysis for the 58 upregulated genes in Jsi1 expressing lines that were also found to be upregulated by *ERF1* and/or *ORA59*. (**C**) GO analysis for the 269 genes found to be enriched in transcription factor binding sites for ERF proteins. Percentage of genes were calculated for each category related to the total number of genes. All GO categories showed *P* values < 0.05. GO categories with more than 20 genes were represented in the graph.

**Figure S8.**
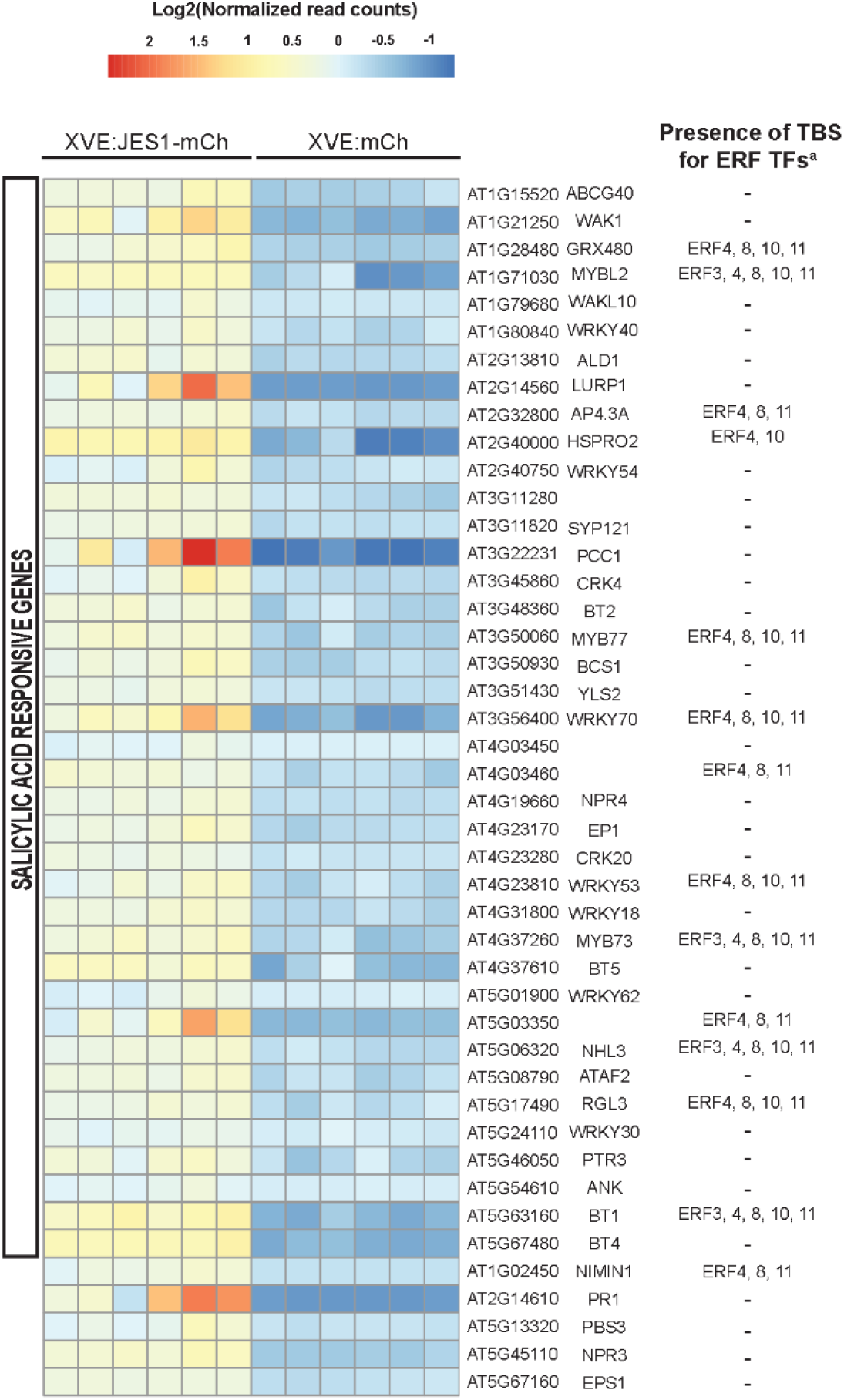
Heat map from RNA-seq showing the GO category for SA responsive genes. *NIMIN1*, *PR1*, *PBS3*, *NPR3* and *EPS1* were included in the heat map as genes related with the SA signaling pathway. a Genes enriched in transcription binding sites (TBS) from ERF TFs with repression activity.

**Figure S9.**
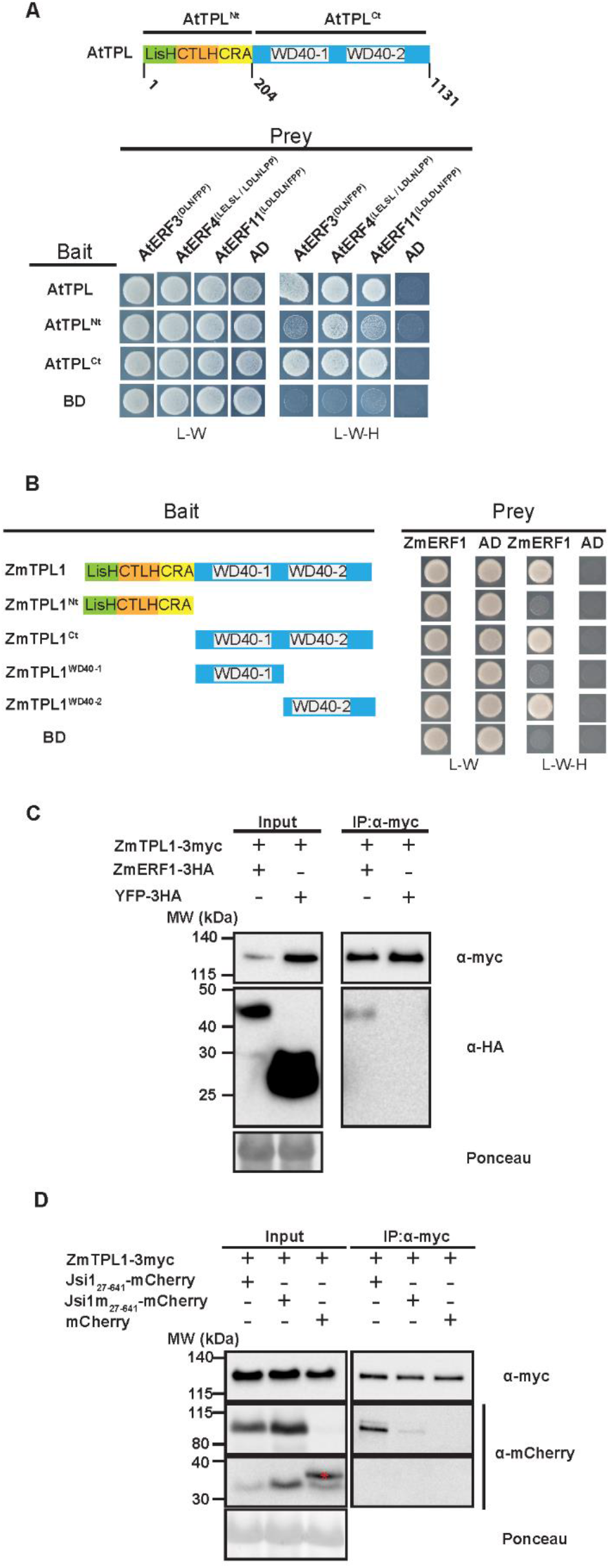
ERF proteins from Arabidopsis and *Z. mays* interact with the C-terminal part of TPL. (**A**) We used AtTPL, N and C-terminal parts of AtTPL as a bait and ERF proteins possessing a DLNxxP motif as a prey to test for interaction in a Y2H assay. (**B**) We tested ZmERF1 containing the DLNxxP motif for interaction with full length ZmTPL1, N and C-terminal parts of ZmTPL1 in Y2H assay. The letters -L, -W and -H indicate medium lacking leucine, tryptophan and histidine, respectively. (**C**) ZmERF1 is in complex with ZmTPL1 *in planta*. We fused ZmERF1 and ZmTPL1 to 3xHA and 3xmyc tag, respectively and co-expressed in *N. benthamiana* leaves. We pulled-down ZmTPL-3myc with anti-myc antibody and the co-immunoprecipitated proteins were detected with anti-myc and anti-HA antibodies. YFP-3xHA was used as a negative control (**D**) Co-IP demonstrates Jsi1 association with ZmTPL1 upon co-expression in *N. benthamiana*. We fused Jsi1 and Jsi1m to mCherry and ZmTPL1 to 3xmyc-tag. We co-expressed Jsi1-mCherry, Jsi1m-mCherry and mCherry constructs with ZmTPL1-3xmyc in *Nicotiana benthamiana* leaves. We pulled-down ZmTPL-3myc and co-immunoprecipitated proteins were detected with anti-myc and anti-mCherry antibodies. Asterisk indicate the mCherry full length protein. mCherry was used as a negative control. Ponceau staining was used as loading control.

**Figure S10.**
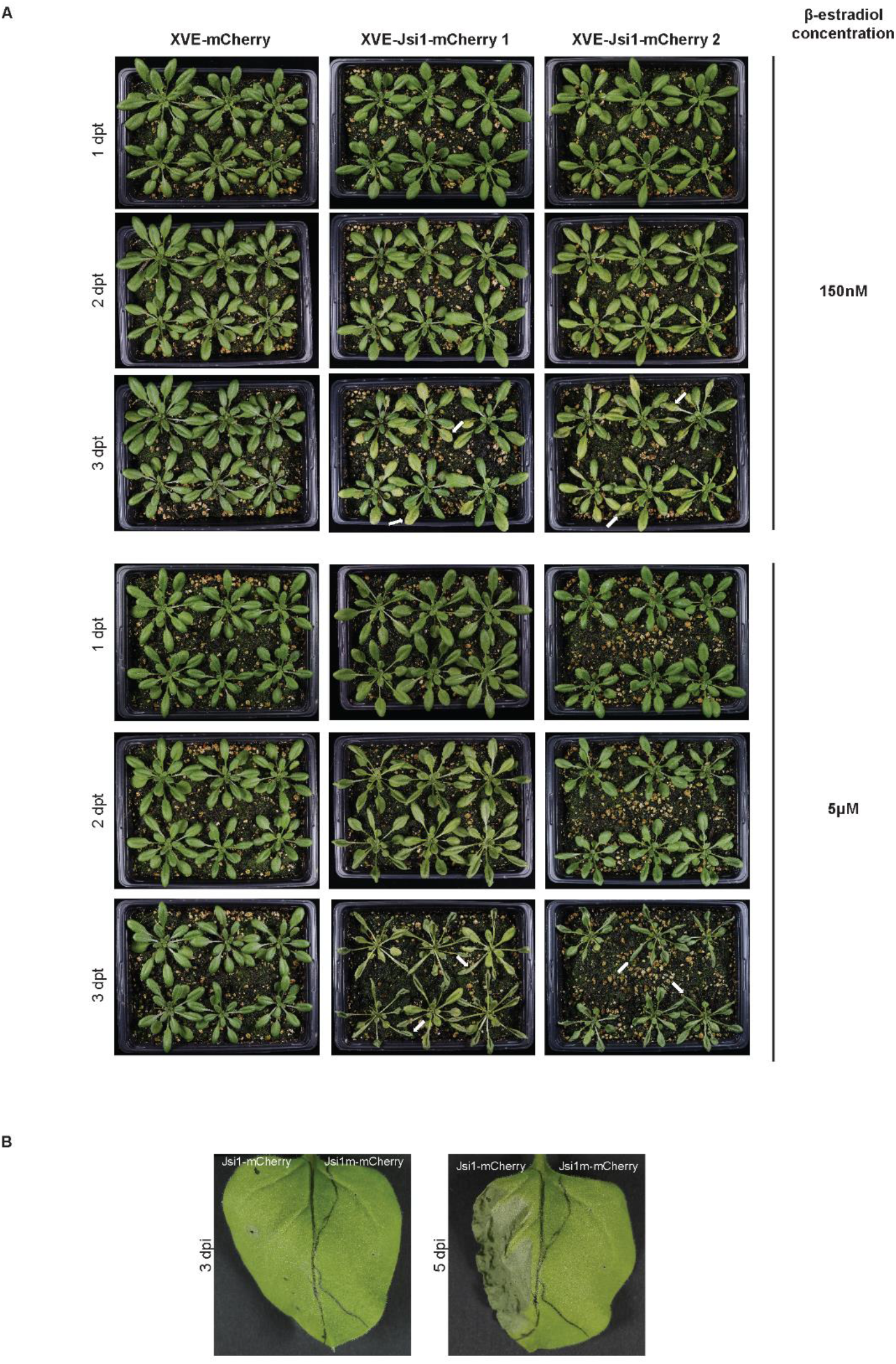
Jsi1 produces cell-death in *N. benthamiana* and Arabidopsis. (**A**) We sprayed 4-weeks old Arabidopsis plants with 150nM and 5μM of β-estradiol. We took pictures at 1-, 2- and 3- days post β-estradiol treatment (dpt). White arrows indicate visible cell-death in Arabidopsis lines expressing Jsi1. (**B**) We infiltrated *N. benthamiana* leaves with constructs expressing Jsi1-mCherry and the mutated version of Jsi1 fused to mCherry (Jsi1m-mCherry) and took pictures at 3 and 5 dpi. Cell-death is visible at 5 dpi only in the leaves side expressing Jsi1-mCherry.

## Supplementary tables footnotes

**Supplementary table 1**. Genes upregulated in Arabidopsis XVE-Jsi1_27-641_-mCherry lines upon Jsi1 induction. ^1^ Genes upregulated in XVE-Jsi1_27-641_-mCherry lines shared with Arabidopsis 35S-*ERF1* line (Lorenzo et al., 2003), and/or Arabidopsis XVE-*ORA59* line2 (Pré et al., 2008). ^3^ We did not detect *PDF1.2* induction in the RNAseq of Jsi1 lines but we detected in the qRT-PCR assay. Fold change and *P values* of genes upregulated by Methyl Jasmonate (MeJa) and/or *Alternaria Brassicicola* (*A. Brassicicola*) were obtained from McGrath et al., (2005).

**Supplementary table 2**. Genes upregulated in Arabidopsis XVE-Jsi1-mCh lines upon Jsi1 expression enriched for ERFs transcription binding sites (TBS). 1 Direct target genes of ERFs were obtained from published DAP-seq data (O’Malley et al., 2016).

**Supplementary table 3.** Constructs used in this study. ^1^ ID number of the constructs. ^2^ pGG187 and pGG446 are plasmids designed for Golden gate cloning using the pGBKT7 and pGADT7 (Clontech®) as backbones respectively. ^3^ pUG is a vector designed from the p123 vector for Golden gate cloning (Stirnberg and Djamei, 2016).

**Supplementary table 4.** List of primer used for RT-PCR analysis. ^1, 2, 3 and 4^, primers pairs from (McGrath et al., 2005, Schellingen et al., 2014, Brown et al., 2003) and (Zhang et al., 2011), respectively.

**Supplementary table 5**. Mass transitions and corresponding conditions for determination of the phytohormones.

